# Understanding the genome-wide transcription response to varying cAMP levels using phenomenological models in bacteria

**DOI:** 10.1101/2022.06.15.496256

**Authors:** Shweta Chakraborty, Parul Singh, Aswin Sai Narain Seshasayee

## Abstract

Attempts to understand gene regulation by global transcription factors (TF) have largely been limited to expression studies under binary conditions of presence and absence of the TF. Studies addressing genome-wide transcriptional responses to changing TF concentration at high resolution are lacking. Here, we create a dataset containing the entire *E.coli* transcriptome as it responds to 10 different cAMP concentrations spanning the biological range. We use the Hill’s model to accurately summarise individual gene responses into 3 intuitively understandable parameters - *k, n* and *Emax* reflecting the midpoint of dynamic range, non-linearity and sensitivity of a gene. cAMP-regulated genes show a small dynamic range with midpoints centred around wild-type cAMP concentrations, with genes activating in a switch-like fashion. Using this approach we show that cAMP-CRP affinity at promoters is well correlated to the sensitivity(*Emax*) of genes but not to the midpoints of dynamic range(*k*). Finally, genes belonging to different functional classes are tuned to different *k, n* and *Emax*. We show phenomenological models to be a better alternative for studying gene expression trends compared to classical clustering methods with the phenomenological constants providing greater insights into how genes are tuned in a regulatory network.

## Introduction

Transcription of a gene is the result of a coordinated effort between transcription factors, various nucleoid-associated proteins and the RNA polymerase at the promoter of a gene. In bacteria, this process is tightly regulated, allowing rapid adaptation and survival in changing environments(1–3). Most studies that focus on understanding the role of a transcription factor in the cell use knock-out strains to study the changes in gene expression in the presence and absence of the transcription factors. The advent of high-throughput RNA sequencing techniques in knock-out strains has accelerated our understanding of the roles of various transcription factors in maintaining the optimum physiology of the cell(4–10). Such experiments give us both qualitative (which genes are differentially expressed) and quantitative information (magnitude of the change).

Concentrations of many transcription factors may exhibit a switch-like behaviour. Their effects may be well modelled by binary states of the transcription factor(11, 12). However, there are many regulators whose concentration in the cell varies continuously (13–16). While studies under binary states of transcription factor concentrations capture snapshots of the gene expression changes, they fail to give any information about the varying response of different genes to changing concentrations of the transcription factor. However, across the genome, different sets of genes in the cell may be tuned to respond differently to changes in transcription factor levels(17–19). In an attempt to better understand the regulatory mechanisms of various transcription factors, previous studies have focussed on gene expression changes in response to modulating transcription factor concentrations(15, 20–23). However, these studies have been limited to a few genes or low resolutions of the regulator concentration. In the recent past, Yang et al (2018), using microarray, have analysed trends in global gene expression patterns at high resolutions of cAMP-CRP activity. Overall, the literature for such studies on a global scale has remained sparse(15, 23, 24).

One example of a transcription regulator whose concentration in the cell varies continuously is the cAMP signalling system in *E.coli*. cAMP is well known for its role in carbon catabolite repression (CCR) and hierarchical utilisation of carbon sources(19, 25–28). cAMP exerts its effect on gene expression by binding and activating the transcription factor CRP (cAMP Receptor Protein)(5, 29, 30). Over years, cAMP along with its effector CRP has been shown to have pleiotropic roles in carbon and nitrogen metabolism, motility, biofilm formation and survival against various stress conditions(31–37). Recent studies have implicated cAMP as a key player in optimising proteome allocation and maintaining flux balance across diverse nutritional conditions(38–40). Intracellular cAMP concentrations are regulated by the nutritional state of the cell. Carbon sources that support high growth rates inhibit cAMP production. As the cell experiences poorer nutritional conditions, intracellular cAMP concentrations increase proportionately. Increasing cAMP levels redirect greater cellular resources towards the expression of transport and metabolic genes(28, 41). *E.coli* cells not only experience cAMP changes across carbon sources(21, 42), but cAMP concentrations in the cell vary continuously in response to cell density and growth phases as well(43–45).

In *E.coli*, the cAMP-CRP complex is responsible for regulating more than 500 genes including close to 100 operons across different conditions(34, 41, 46). Since cells experience a continuous change in cAMP concentrations, we wanted to quantitatively characterise the different trends each gene may follow in response to activation by cAMP. Previously, Liu et al have qualitatively described the response of cAMP regulated genes to increasing cAMP concentrations achieved by altering carbon sources(21). Other works have shown that it is possible to control activities of cAMP regulated genes by modulating extracellular cAMP, allowing easy experimental control over this system(22, 24, 33, 47, 48).

In this study, we expose cAMP deficient *ΔcyaA E.coli* mutants to 10 different concentrations of extracellular cAMP and measure the global gene expression changes in response to varying cAMP concentrations using RNA-Seq. We use phenomenological models to quantitatively describe the different trends that individual genes followed in response to increasing levels of cAMP. We show that most cAMP responsive genes follow a sigmoid curve which can be satisfactorily modelled using Hill’s equation. Presence of a sigmoid type response curve in gene expression circuits is not uncommon. As a matter of fact, the prevalence of feedback and feedforward loops in regulatory networks make the sigmoid response motif extremely common. Further, the use of the Hill class of equations to model dose-response curves has been found to be quite useful in understanding interaction kinetics in biochemical and pharmacological studies. In the recent past, it has been found to be useful in the modelling of transcription regulatory networks as well(49–54).

In the next part of the paper, we try and explore the use of the phenomenological constants obtained from Hill’s model to gain insights into the workings of the cAMP regulatory network. In this study, we use the 4 parameter Hills model defined by *b0, n, k* and *Emax*. The biological implications of these constants have been hashed out for single transcription factor – gene interactions(50, 52, 55–57). We attempt to use these parameters to study the properties of the cAMP regulatory network.

## Results

### 1. Effects of increasing cAMP concentrations on growth rates of *ΔcyaA E.coli* mutant

Adenylate cyclase is encoded by the *cyaA* gene in *E.coli* and converts ATP to cAMP. *ΔcyaA E.coli* mutants cannot make intracellular cAMP(29, 58). *E.coli* cells that lack the *cyaA* gene are capable of growing in rich permissive media like LB, albeit with a growth defect (S1A, B). However, these mutants exhibit a complete loss of growth when growing in sugars like lactose, sorbitol, ribose etc. that require the cAMP-CRP signalling system for their utilisation (S1C, D).

We wanted to see if the addition of extracellular cAMP can rescue the growth defect induced by *cyaA* deletion. To this end, we studied the growth dynamics of *ΔcyaA* cells across 10 different extracellular cAMP concentrations ranging from 0mM to 4mM cAMP in 3 different media – LB, M9 with lactose and M9 with mixture of sorbitol and ribose. To determine if the extracellular cAMP enters the cell, we measured the levels of intracellular cAMP at each administered dose. We found a linear relationship between the measured intracellular cAMP and extracellular cAMP provided(S1G). For the rest of the study, all instances of cAMP refer to the extracellular concentrations provided, unless mentioned otherwise.

As expected, *ΔcyaA* (0mM cAMP) cells showed a lower growth rate in LB compared to the wild-type (WT) cells (Fig1). In M9 media with sugar, *ΔcyaA* cells failed to grow in the absence of cAMP (S1C, D). Irrespective of the media, exposure to varying concentrations of cAMP induced an increase in growth rates in a monotonic fashion, with growth rates being restored to wild-type levels at high cAMP concentrations, between 0.8mM – 4mM cAMP (Fig1, S1C, D). In LB, growth rates reached wild-type levels between 0.6-0.8mM cAMP. We measured the concentration of cAMP in wild-type *E.coli* cells growing in LB to be 5.4(±1.06) pmol/OD/ml (S1B). It is equivalent to 0.72 (±0.12) mM extracellular cAMP, which is close to the administered cAMP concentration at which the *ΔcyaA* cells reached wild-type levels. We observed that no change in growth rates was induced till 0.3mM cAMP. This could be because the amount of cAMP entering the cells at low concentrations was insufficient to effect a transcriptional response.

**Figure1.**
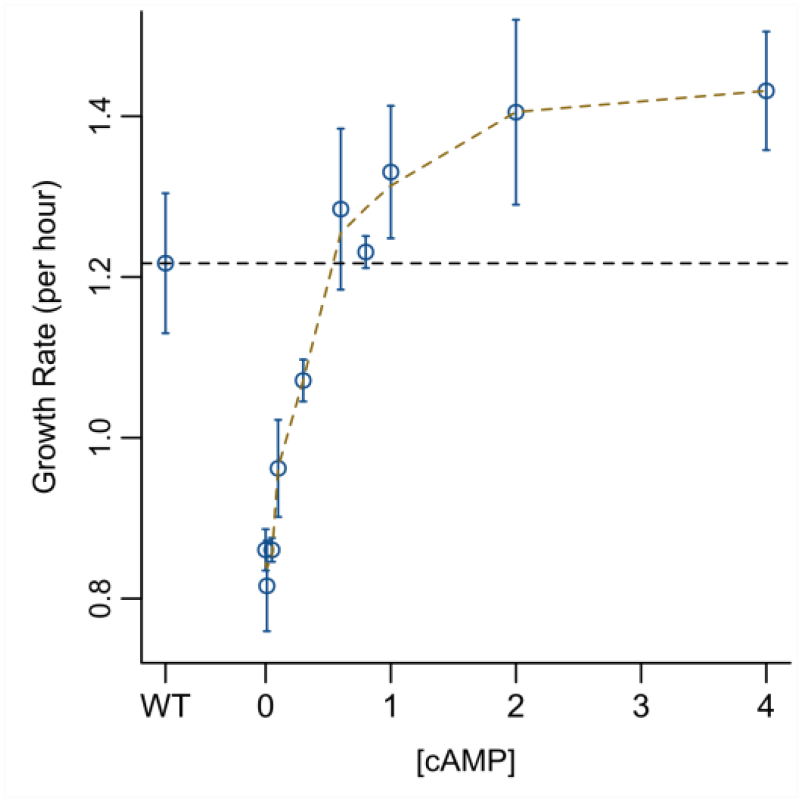
Changes in growth rate with increasing cAMP dosage. Growth rates of wild-type and *ΔcyaA E.coli* cells in LB media with increasing doses of cAMP. 0mM cAMP represents the *ΔcyaA* strain. Growth rates of *ΔcyaA* cells increase monotonically in response to increasing extracellular cAMP concentrations. Each point represents the mean of 3 replicates with standard deviation. Growth curves for the wild-type and *ΔcyaA* mutant growing in LB are shown in S1A.

One difference in growth dynamics for cells growing in M9 minimal media with sugars compared to LB was in their lag-times. In batch culture, lag-time refers to the time taken by a population to exit the lag phase and transit to the log phase of growth. In LB, the *ΔcyaA* cells showed no change in lag-time in response to the cAMP concentration gradient. However, in lactose and sorbitol-ribose media cells experienced decreasing lengths of lag-time with increasing cAMP concentrations(S1E-F).

### 2. Global transcriptional changes in *ΔcyaA* mutant in response to ordered exposure of cAMP

The cAMP-CRP signalling system in *E.coli* is a global regulator of transcription. Since the addition of extracellular cAMP was able to restore the growth rates of *ΔcyaA* mutant to wild-type levels, we wanted to see if it had the same effect on its transcriptome as well. We grew *ΔcyaA* cells in LB with 10 increasing doses of extracellular cAMP followed by whole transcriptome RNA sequencing at each cAMP concentration.

cAMP and CRP have been reported to regulate more than 500 genes in *E.coli* across various conditions (5, 6, 41). In LB with no added cAMP, we found 488 genes to be differentially expressed between wild-type and the *ΔcyaA* mutant in LB (logFC > ±1 and p-value < 0.01). 305 genes were upregulated in the wild-type strain compared to the *ΔcyaA* mutant while 183 genes were down-regulated. For any given gene, we calculated its response to a cAMP concentration as the fold-change in expression experienced by the gene at that cAMP concentration compared to the 0mM cAMP state (*ΔcyaA*). We observed that genes under cAMP control responded monotonically to increasing concentrations of extracellular cAMP with positively regulated genes expressing and negatively regulated genes repressing monotonically with increasing levels of cAMP (Fig2A, B). Since cAMP-CRP is primarily a transcriptional activator, in this study we concentrated only on genes which are under positive regulation by cAMP.

**Figure2.**
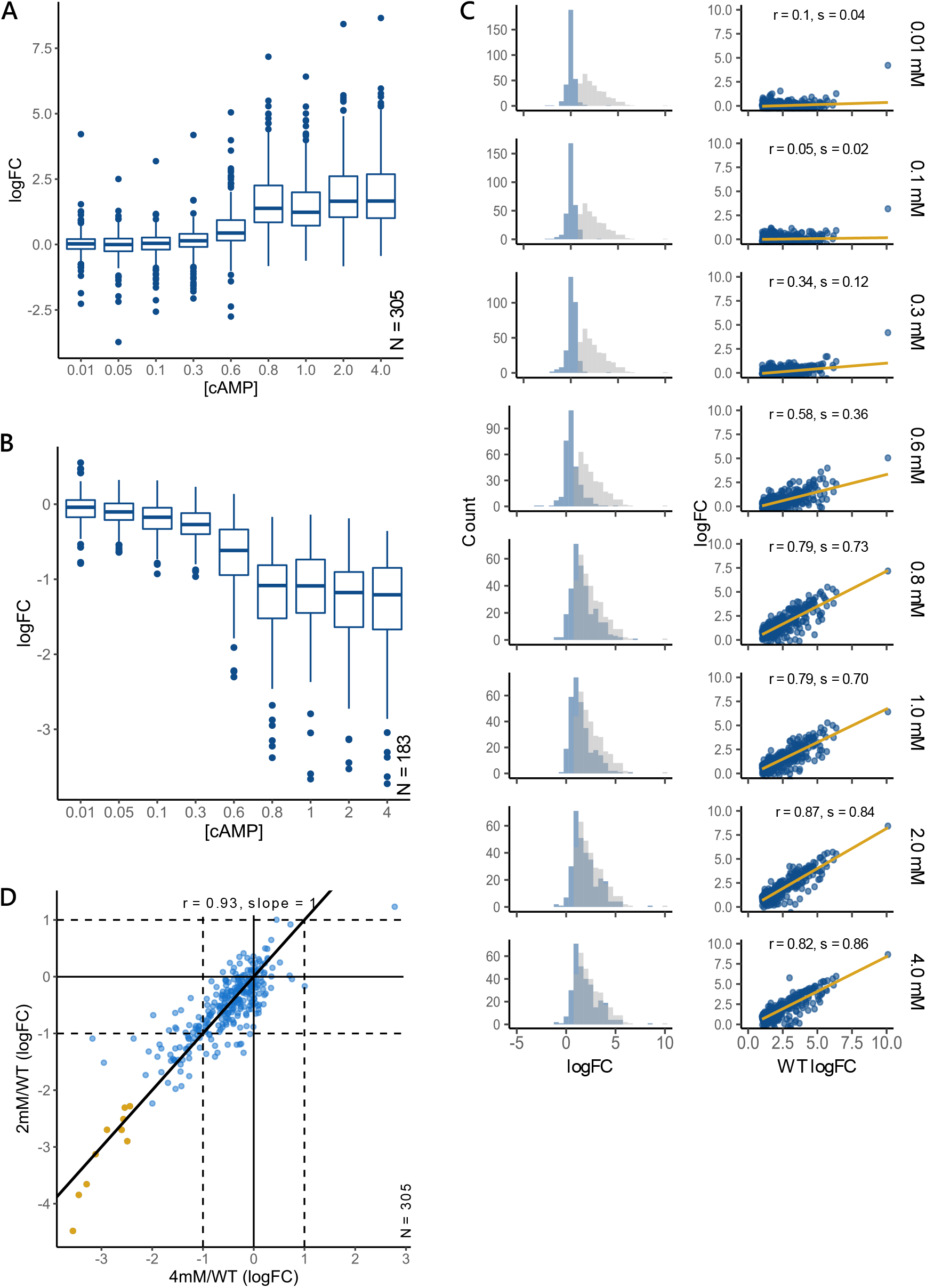
Expression changes of the cAMP regulon in response to increasing cAMP concentrations. **(A-B)** Distribution of expression values at different doses of extracellular cAMP for genes under positive (A) and negative(B) regulation of cAMP. In LB, 305 genes are positively and 183 genes are negatively regulated by cAMP in the wild-type strain. For a gene, change in expression was calculated as the log foldchange at a cAMP concentration compared to the 0mM cAMP *ΔcyaA E.coli* strain. **(C)** Left panel shows the frequency distribution of gene expression at each cAMP concentration(blue) compared to the wild-type distribution (grey). Right panel shows the correlation between gene expression at each cAMP concentration with wild-type gene expression.**(D)** Scatterplot for correlation between gene expression at 2mM and 4mM cAMP concentrations. Each point represents a gene. Strong Pearson correlation of 0.93 (*p-value* < 0.01) show that gene expression has saturated by 2mM cAMP

Consistent with our observations in the growth rate studies in LB, we observed changes in gene expression only post 0.3mM cAMP, followed by a monotonic increase in expression (Fig2C). A large number of genes expressed between 0.6mM and 0.8mM cAMP, beyond which changes in gene expression started saturating(S2C). We observed that 0.8mM cAMP was sufficient to induce wild-type levels of growth rates in *ΔcyaA* cells (Fig1) but not sufficient to restore gene expression levels to that of wild-type (Pearson correlation, r = 0.79, slope = 0.73, *p-value* < 0.01; Fig2C). Despite many genes failing to reach wild-type, gene expression levels at 0.8mM cAMP were sufficient to gain wild-type growth rates, indicating that all differentially expressed genes did not contribute equally towards the maintenance of cellular growth rates.

A strong correlation of 0.93 with slope of 1 (p-value < 0.01) between gene expressions of 2mM cAMP and 4mM cAMP revealed that expressions saturated by 2mM cAMP (Fig2D). By 2mM cAMP, the transcriptome is also well correlated with that of the wild-type (Pearson correlation, r = 0.82 and slope = 0.84, *p-value* < 0.01). However, even at high concentrations of extracellular cAMP, not all cAMP regulated genes reached wild-type levels of expression with the expression of many genes remaining lower than those of the wild-type (Fig2D yellow points). Despite administering doses of cAMP 4 times that of the wild-type concentrations, only 10 genes were differentially expressed between the 4mM cAMP and wild-type(S2B).

The cAMP-CRP complex binds to promoters and recruits RNA polymerase (RNAP) facilitating the transcription of a gene. We wanted to study the effects of CRP or RNAP binding at the promoter on gene expression in the wild-type cell. We used signals obtained from chromatin immunoprecipitation (ChIP) studies to calculate the occupancy of CRP or RNA polymerase at a promoter. For CRP, we used ChIP-seq to measure the CRP occupancy across the entire chromosome in exponentially growing wild-type cells. For RNAP, publicly available ChIP data(59) was used to calculate the RNAP occupancy at the promoter of each gene or operon(Methods 4). The median of the distribution for both CRP and RNAP occupancy at promoters of genes that are positively regulated by cAMP was significantly higher than that of non-differentially expressed genes (Wilcoxon Rank Sum test, *p-value* < 0.001; S3A-B). Further, we found a significant correlation of 0.46 (*p-value* = 4.14 × 10^−12^) between the foldchange in expression (wild-type with respect to *ΔcyaA* mutant) of genes positively regulated by cAMP and their CRP occupancy. However, no such correlation was observed for RNAP occupancy and expression(S3C-D).

Given that cAMP activated close to 300 genes in LB, studying variations in individual gene expression kinetics in response to cAMP can give deeper insights into the nature of the cAMP-CRP regulatory network. To this end, we attempted to characterise the various trends that genes may follow in response to the ordered exposure of cAMP.

### 3. Use of phenomenological models to describe cAMP regulated genes

In the previous section, we quantified changes in gene expression induced by cAMP exposure. To generate dose-response curves for each gene, we calculated the effect of a cAMP concentration on a gene as the change in expression (foldchange) of the gene at that cAMP concentration relative to its expression in 0mM cAMP *ΔcyaA* mutant. To classify and quantitatively summarise these dose-response curves, we made use of phenomenological models. Phenomenological models are a useful tool to quantitatively describe trends independent of the underlying mechanisms that produced them. Further, the phenomenological constants obtained from these models can be used for the characterisation of the observed behaviour.

Dose-response curves often follow a sigmoid behaviour. Such non-linear behaviour of gene expression in response to activation by transcription factors is well known in biological systems. The presence of feedforward loops in regulatory networks and cooperative behaviour of transcription factors in enhancing their own binding at the promoter leads to sigmoid behaviour(60–62, 56). Hill-type models best describe such sigmoid trends. We used the four parameter Hill’s model, defined by *b0, k, n* and *Emax*, to characterise genes following sigmoid behaviour in our data(50). It is possible that some genes do not reach their saturation levels under the regime of the experiment. These would appear to have a non-saturating trend. We used the linear model to describe such a response. We compared these models against a null model defined by no change in response to the given signal (Methods 5).

Overall, we used 3 models to bin the dose-response curves – the null model (NR) to identify genes that show no change in expression in response to increasing cAMP concentrations, the Hills model (HM) to fit genes which show sigmoid and first order Michael – Menten behaviour and the Linear Model (LM) for genes that exhibit non-saturating behaviour within the cAMP range used in this study. Gene trends satisfied Hill’s model (HM) only if the residual sum error (σ^2^) of the fit by HM was less than that of the competing models, its R^2^ > 0.8 (*p-value* < 0.05) and slope = 1±0.2 and relative standard error (RSE) for predicted parameters *k* and *Emax* < 20%. Since the number of data points was less around the transition points, the predictions for *n* showed a larger variation in RSE. Hence, we did not use the RSE of *n* as a cut-off factor. Similarly, we considered genes to fit the linear model (LM) if σ^2^ for the fit was less than other models, its R^2^ > 0.7 and *p-value* < 0.05. Genes followed the null model if the σ^2^ for the fit was less than the other models and R^2^ > 0.7 and *p-value* < 0.05 (Methods 5(b-c)).

Using the model fit method described above, we found that 74% (225/305) of cAMP regulated genes followed the HM, while 9% (28/305) genes followed LM. Despite being differentially expressed in the wild-type strain compared to the *ΔcyaA* mutant, 6% (18/305) genes fit the null model (NR) and did not show any change in expression in response to the extracellular cAMP. 11% (34/305) of genes did not fit any of the above models. These genes showed a non-monotonic (NM) response to increasing cAMP concentrations (Fig3A).

**Figure3.**
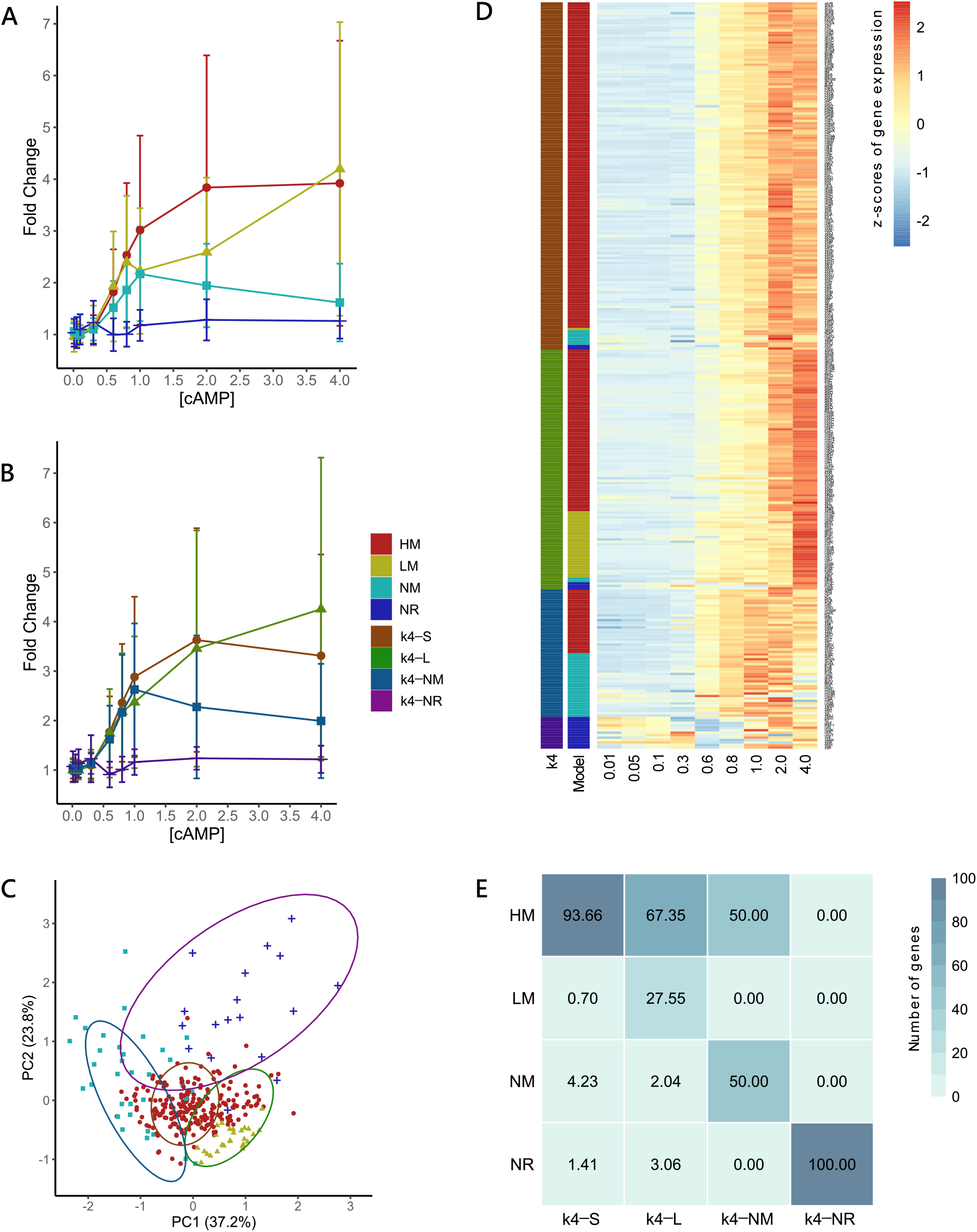
Comparison of different clustering methods. **(A-B)** Median trends for clusters from the model-fit method (A) and k-means with 4 clusters (B). Error bars represent the median absolute deviation (MAD) around the median for each cluster at different cAMP concentrations. Broadly, cAMP regulated genes follow one of 4 trends – Sigmoid(HM/k4-S), linear(LM/k4-L), non-monotonic(NM/k4-NM) or non-responsive(NR/k4-NR).**(C)** Projection of gene expression trends on a PCA plot. Each point represents a gene. Different shapes indicate the cluster a gene belongs to using the model-fit method. Ellipses indicate the gene clusters formed using the k-means clustering methods.**(D)** Comparison of genes across clusters for the k-means and model-fit method. Each row represents a gene. For the two clustering methods, colours represent the cluster a gene belongs to. The heatmap shows the normalised expression levels for the gene at each cAMP concentration.**(E)** Correlation plot between genes binned by the model fit and k-means with 4 clusters. Numbers in the boxes represent the percentage of k4-X genes that follow the mode-fit trends.

Unsupervised clustering is one of the common methods used to study trends in a data set(63–65). We compared the model fit method described above to the results obtained using clustering algorithms like the *k*-means and hierarchical. Both these methods revealed the continuous nature of the data and the lack of clear clusters in it. This was reflected in the inability of methods meant to find optimum clusters for *k*-means to reach a consensus (S4A) and the presence of large cross-correlation across clusters in the distance matrix (S4E). Based on the Silhouette method, we divided the genes into 3 clusters using *k*-means. The *k*-means algorithm yielded 2 major clusters containing 31.1% (95/305) and 64.2% (196/305) of the genes and a small cluster with 4% (14/305) genes. The two major clusters showed a median trend resembling sigmoid curves(S4B-C).

Since the model fit method showed that cAMP regulated genes fall into 4 clusters, we forced the *k*-means algorithm to divide the data into 4 clusters. This yielded patterns similar to those from the model fit (Fig3B). For easier visualisation, we overlaid the genes into groups partitioned by *k*-means and model fit on 2D PCA plot (Fig3C). We observed that gene trends are in a continuum with partitioning by both methods happening primarily across the first principal component (PC1) where gene trends vary from NM to HM to LM. Each *k*-mean cluster primarily corresponded to at least one trend from the model fit (Fig3C-E). Clusters generated by the *k*-means algorithm largely agreed with partitions made by the average linkage hierarchical clustering method used to analyse the gene trends (FigS5).

In this section we showed that genes under the positive regulation of cAMP broadly follow 4 trends – Sigmoid, linear, non-monotonic and non-responsive. The overall gene expression response to cAMP populates a continuum as compared to distinct clusters, with majority of the genes following a sigmoid dose response curve that could be described quantitatively by phenomenological models like the Hill’s. We note that conventional clustering methods can in fact meaningfully partition the data into intuitively understandable shapes. However, finding the optimum clusters in a continuous data poses a difficult decision. Model fitting circumvents this problem by adding another layer of clarity, making it possible to quantitate the variations observed across these continuous trends. Not only this, parameters obtained from the Hill’s model can be used to further characterise the behaviour of these dose response curves.

### 4. Characterisation of phenomenological constants obtained from the Hill’s model

For a gene whose transcription is activated by cAMP, the Hill’s model defines a sigmoid dose response curve using 4 parameters: (1) *b0*, represents the basal level of gene expression when no cAMP signal is observed, (2) *k*, quantifies the levels of cAMP required for the gene to reach half its saturating concentration, (3) *Emax* quantifies the magnitude of foldchange at the saturating concentration and (4) *n* describes the rate of change of expression in response to changing cAMP levels(Fig4A).

**Figure4.**
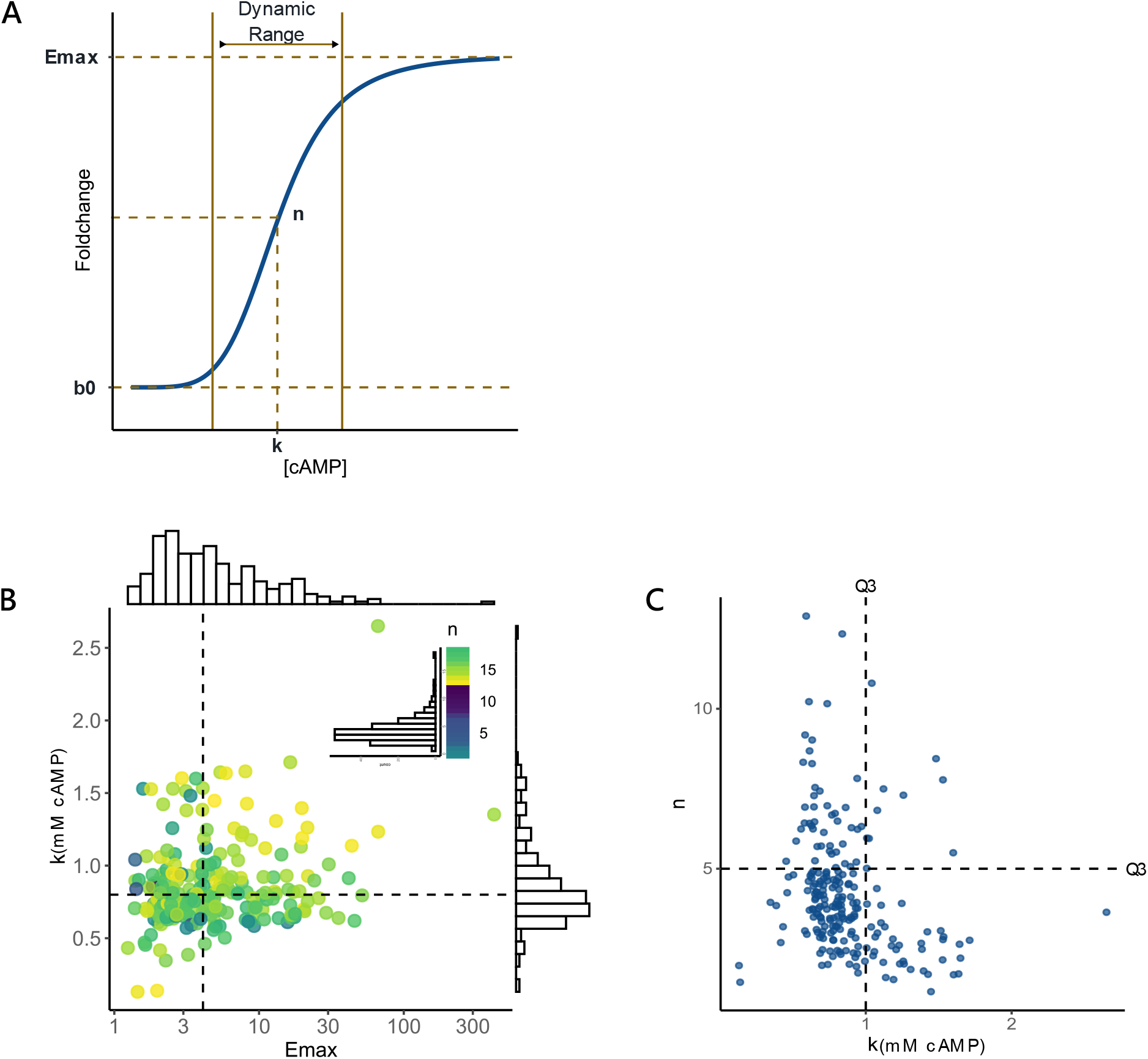
Distribution of the parameters obtained from the Hill’s model. **(A)** Schematic showing the dose-response curve for a gene ollowing the Hill’s model, with the 4 parameters – *b0, k, n* and *Emax* marked. *k* and *n* together determine the dynamic range of a gene. **(B)** Distribution of *k* and *Emax* or genes that ollow the Hill’s model he colour o each point represents the value of *n*. There is no correlation between *k* and *Emax* or *n* and *Emax* (data not shown). Dashed lines show the median of the distributions of *k* and *Emax* **(C)** Scatterplot showing the absence of high *k* and high *n* genes in the study. Dotted lines represent the value of top third quartile(Q3) for the respective distributions. N = 225

Biologically, cAMP concentrations at *k* reflect the midpoint of the dynamic range of a gene. It determines when in the cAMP concentration a gene responds. Except 2 genes, all genes showed a *k* greater than 0.35mM, with values of *k* ranging up to 2.6mM cAMP (Fig4B). This was consistent with the findings from global gene expression analysis where we observed no change in gene expression prior to 0.3mM cAMP (Fig1). Genes under cAMP control showed a median *k* of 0.79mM cAMP with an interquartile range (IQR) of 0.26mM. The IQR of *k* accounted for only 20% of the entire range of *k*, implying that the mid-point of dynamic range of many cAMP regulated genes (i.e., 50% genes) occupy a very small space in the range of *k*. Further, we observed that the median value of *k* lay close to the wild-type levels of cAMP for *E.coli* growing in LB (0.72mM cAMP). The observation that the midpoints of the dynamic range of gene expressions (median *k*) is centred around the wild-type concentration of cAMP suggests that promoters and intracellular cAMP concentrations in the cAMP-CRP regulatory network may be tuned to have optimised gene expressions.

*Emax* defines the maximum expression of a gene when cAMP is not the limiting factor. It reflects the sensitivity of a gene to cAMP. We observed that the distribution of *Emax* for cAMP regulated genes in LB is right skewed with saturating gene expressions varying from 1.2 to 66.5 fold change (except *gatB* which showed a very high *Emax* of 420.7 foldchange). The variation observed in *Emax* (MADM = 48.4%) was much greater than that of *k* (MADM = 16.5%), showing that differences in the *Emax* of genes affected the observed inter-gene variation in expression more than the differences in *k*(Fig4B).

*n* determines the steepness or non-linearity of the dose-response curve. Non-linearity in the system is introduced by the cooperative behaviour of transcription factors and the presence of positive feedforward and feedback loops. Higher *n* indicates more switch-like behaviour while low *n* indicates a more graded response. All genes following the HM exhibited a *n* > 1. *n* for cAMP regulated genes showed a median value of 3.9 with an IQR of 2(S6B). The high values of *n* for cAMP regulated genes reflect the pervasive nature of feedback and feedforward loops in the network resulting in switch-like behaviour for most genes in the population. For the use of this study, we considered values of *n* > 5 to be high, between 3-5 moderate and *n* < 3 low.

We observed a lack of genes that had both high *n* (>5) and high *k* (> 1) in the population (Fig4C). Very few genes (like *torR, glpK, ybhG, ilvX*) showed both high *n* and *k*(S6D). This led to an apparent negative relationship between the *n* and *k*. Hence, genes which saturated at high cAMP concentrations (high *k*) were likely to behave in a relatively graded manner (have low *n*) in response to cAMP. Conversely, genes which showed a more switch like behaviour (high *n*) were more likely to also saturate at lower cAMP levels (have low *k*). This pattern could also be the result of the experimental design, as it may be difficult for the algorithm fitting the Hill’s model to capture genes with trends having high *k* and *n* due to the lack of data points depicting the transition and saturation states at high cAMP concentrations. Although upon manually checking the trends of genes rejected by the Hill’s model, we noticed that no gene showed such a trend, hinting that the effect may result from biological limits as opposed to experimental limitations.

Overall, our data showed that expression of cAMP regulated genes in LB differed largely in their *Emax* compared to *k*. Also, instead of having a graded response across the biological range of cAMP, most cAMP regulated genes exhibited a switch like behaviour with their dynamic range centred around 0.79mM cAMP, close to the wild-type concentrations of cAMP in LB. This resulted in genes rapidly switching on and reaching saturation (*Emax*) at concentrations close to 0.79mM cAMP. This is consistent with the observation that gene expression increased rapidly between 0.6 and 0.8mM cAMP (Fig2C; S2C).

### 5. Comparison of Hill’s parameters across genes under direct and indirect control of cAMP

The cAMP regulatory network consists of genes that are directly regulated by the cAMP-CRP complex binding at their promoter as well as genes which are under indirect control via intermediate players. We binned the cAMP regulated genes as direct or indirect targets of the cAMP-CRP complex using 2 different criteria. In the first classification method, we considered genes to be direct targets of the cAMP-CRP complex if they were also annotated to be directly regulated by CRP in the Ecocyc/RegulonDB database (Direct_Ecocyc_). We binned the rest of the genes as Indirect_Ecocyc_. 66/225 HM genes belonged to Direct_Ecocyc_. Overall, 98/305 genes belonged to Direct_Ecocyc_. The Direct_Ecocyc_ genes showed a greater CRP occupancy compared to Indirect_Ecocyc_ genes (Wilcoxon Rank sum test, *p-value* = 0.0092; S7A), confirming that the set of genes are indeed under direct regulation of cAMP. However, we noticed that genes belonging to Indirect_Ecocyc_ showed CRP occupancy scores greater than those of non-differentially expressed genes(S7A). Since the assumption is that the cAMP-CRP complex only binds to promoters of direct genes, one would expect genes under indirect regulation of CRP to have occupancies similar to those of non-differentially expressed genes. This indicated that the Ecocyc/RegulonDB defined set of direct genes may not be complete.

We defined a more inclusive set of Direct_ChIP_ and Indirect_ChIP_ genes based on the data obtained from Ecocyc/RegulonDB database and CRP occupancy scores obtained from the ChIP experiments. The set Direct_ChIP_ was defined by taking the union of Direct_Ecocyc_ and those genes in Indirect_Ecocyc_ that had a CRP occupancy score greater than 2.8. This cut off was chosen based on the bottom quartile(Q1) of CRP occupancy scores of the Direct_Ecocyc_ set(S7B). 147/225 HM genes belonged to the Direct_ChIP_ set. Of all cAMP activated genes, 211/305 were binned as Direct_ChIP_ (S7C).

For an activator, *k* reflects the transcription factor concentration around which the gene behaves most dynamically for cases where n>1. For a gene that is under direct control of cAMP, *k* is affected by the affinity of the transcription factor to the promoter. For indirect genes it reflects the composite effects of all *k*s in the network. *n* determines the steepness of the response curve and is affected by the presence of multiple regulators, feedforward and feedback loops present in the circuit. Together, they determine the dynamic range of a gene in response to the transcription factor. Thus, the observed *k* and *n* of a gene in the regulatory network may get affected by its level of regulation. Since we did not have any formal expectation of how these parameters may be affected by their levels of regulation, we used a toy model of a simple regulatory circuit to determine how the distributions for these parameters should look across direct and indirect genes.

We considered a linear regulatory chain having 4 nodes and 3 edges, with each edge having independent *k* and following the Hill’s function (Fig5A). *k* at each level was chosen randomly from the distribution of observed *k* from our study. The first edge is akin to the regulation of a direct gene by cAMP. The second edge takes concentrations of the direct gene(g1) as an input function for the first indirect gene (g2, indirect level1). We mapped the outcome at each node to the input concentrations of cAMP and calculated the apparent *n* and *k* at each level. For this simulation, we did not consider changes in *Emax. Emax* reflects the maximum foldchange a gene can experience when cAMP is unlimited. It is more likely to be affected by promoter related properties (described in later sections) than the level of regulation. Thus, for simplicity, *Emax* was set to 1 for the toy model. Since we found all HM genes to have n >1, in this circuit, we fixed the values of *n* at 2 and *Emax* at 1. The mean and variance for distributions of apparent *k* and *n* increased at each regulatory level (Fig5B, C). Thus, our null model predicted that the response of genes further down the regulatory circuit show (1) greater non-linearity(*n*) and (2) larger variation in their dynamic range(*k*).

**Figure5.**
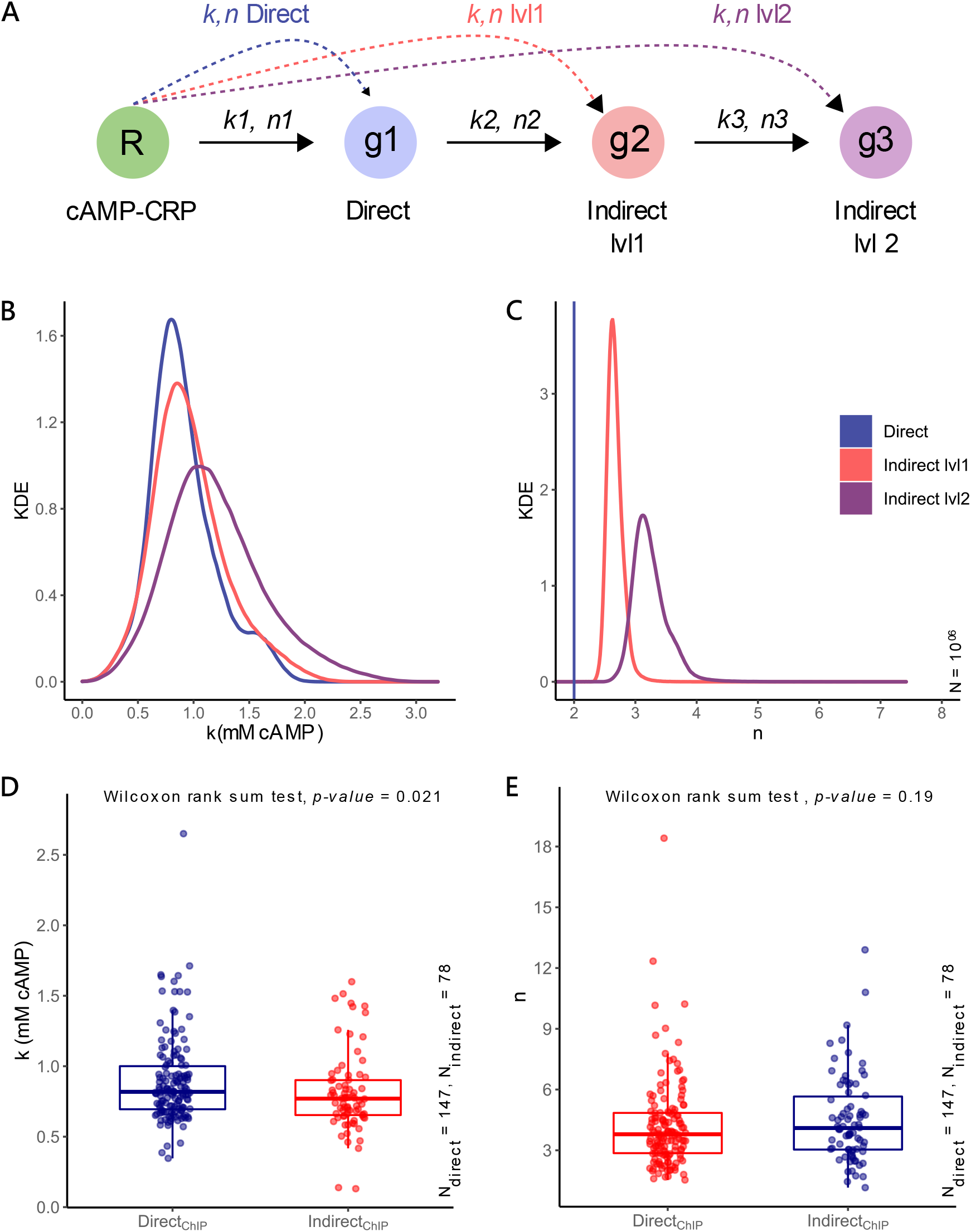
Comparison of k and n across direct and indirect genes. **(A)** Toy model of a linear transcription network representing a regulator (R), a directly regulated gene (g1) and two indirectly regulated genes (g2 and g Each edge ollows the Hill’s unction *k* at each edge is independent of other edges and is chosen at random from the same distribution. *Emax* and *n* remain constant across the circuit, with *n1*=*n2*=*n3* =2 and *Emax* = 1. We calculate the apparent *k* and *n* of the response for g1(direct), g2(indirect lvl1) and g3(indirect lvl2) with respect to the regulator(R). (**B-C)** Expected distribution of apparent *k* and *n* at each level of the regulatory network for 10^06^ trials. **(D-E)** Observed distributions of *k* and *n* across Direct_ChIP_ and Indirect_ChIP_ sets of cAMP regulated genes. We used Wilcoxon rank sum test to compare the distributions.

Contrary to the null model, our data showed no differences in the distribution of *k* and *n* across direct and indirect genes, irrespective of the definition of direct and indirect genes used (Wilcoxon rank sum test, *p-value* > 0.01, Fig5D-E; S7E-F). Deviation from the null model reflected that either one or both assumptions (linear topology and independent *k* at each level) of the simple model were violated. We suspect that the presence of feedback loops feeding into higher nodes could cause the variation in *n* of genes to increase, leading to comparable distributions of *n* for direct and indirect genes. This finding implies that direct genes are not regulated solely by cAMP and that cyclic edges are pervasive even among direct genes. Presence of cyclic edges feeding into higher nodes could also cause the *k*s at each step to be dependent on each other, leading to violation of the null model. Another reason could be that the cAMP regulatory network mostly consists of short length subnetworks limiting the variation in *k* for indirect genes.

### 6. Contribution of CRP and RNA polymerase binding on gene expression

A successful transcription event is the product of concerted interactions of various molecular players like regulators and helper NAPs with the RNA polymerase (RNAP) at the promoter. Models like the Hill’s can help abstract the strength of these interactions into physiologically relevant phenomenological constants. For cAMP regulated genes, two major players that affect the gene expression are the cAMP-CRP complex and RNA polymerase. As mentioned above, *k* for direct binding genes reflects the affinity of the cAMP-CRP complex to that of the promoter while *Emax* is affected by promoter properties like RNAP binding and CRP binding to the promoter. Since *k* and *Emax* may capture the physiological effects of the cAMP-CRP and RNAP binding to the promoter on gene expression, we asked if their binding strengths have any effect on the variation of these parameters across cAMP regulated genes. We used data from ChIP experiments in *E.coli* as a proxy for binding affinities of CRP and RNAP. We assumed the strength of the signal at a position to be proportional to the probability of the target molecule occupying that position, which in turn is affected by their effective binding affinity.

We checked the correlation between *k* and cAMP-CRP occupancy scores for Direct_ChIP_ genes. *k* of cAMP regulated genes did not show any correlation with cAMP-CRP occupancy (Fig6A; S8A-B), indicating that CRP occupancy alone was not sufficient to explain the observed values of *k*. These results hinted that for a cell growing in a specific medium, individual binding affinities may not play a large role in determining the midpoint of dynamic range of genes.

**Figure6.**
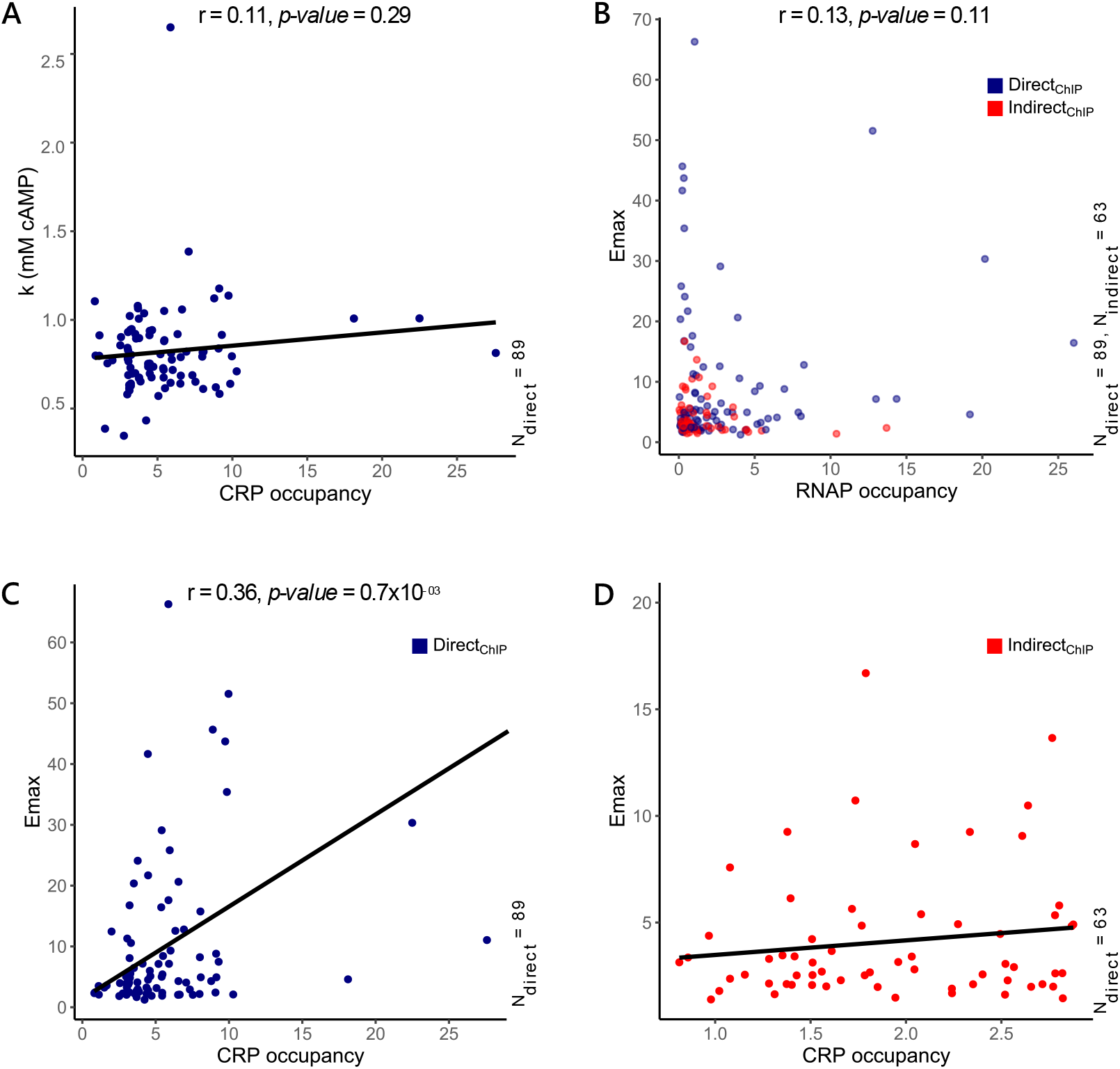
Effects of CRP and RNAP occupancy on *k* and *Emax* of cAMP regulated genes. **(A)** Correlation between *k* and CRP occupancy for genes under direct regulation of the cAMP-CRP complex(Direct_ChIP_). **(B)** Correlation between *Emax* and RNAP occupancy for both Direct_ChIP_ and Indirect_ChIP_ genes. **(C-D)** Correlation between *Emax* and CRP occupancy for Direct_ChIP_(C) and Indirect_ChIP_ genes(D). r and *p-value* in the plots represents the Pearson correlation coefficient and the corresponding *p-value* for the given pair of variables.

*Emax* of a gene reflects the maximum change in expression(foldchange) it can experience when cAMP is not limiting, i.e., [cAMP] >>*k*. It reflects the maximum sensitivity of a promoter to cAMP. *Emax* is affected by promoter properties like RNAP binding affinity, interaction of CRP, other regulators and different NAPs with the RNAP, promoter escape rates and gene dosage(66–68). We observed no correlation between *Emax* of cAMP regulated genes with RNAP occupancy at the promoter (Pearson correlation, r = 0.13, *p-value* = 0.11, Fig6B). CRP occupancy, on the other hand, showed a small but significant correlation of 0.36 (Pearson correlation, *p-value* = 10^−06^) with *Emax* (Fig6C-D; S8C-D). We observed that genes belonging to Direct_ChIP_ had a significant correlation between *Emax* and CRP occupancy (Pearson correlation coefficient r = 0.36, *p-value* = 0.7 × 10^−03^) while Indirect_ChIP_ genes did not show any correlations (Pearson correlation coefficient, r = 0.13, *p-value* = 0.29), proving that the observed correlation between *Emax* and CRP occupancy for cAMP activated genes was driven by directly binding genes.

In this section, we have attempted to quantify the individual effects of two major molecular players -CRP and RNAP on the dynamic range(*k)* and sensitivity (*Emax)* of genes under direct control of cAMP. Our data showed that despite having different binding affinities to the promoters, under physiological conditions cAMP-CRP binding affinities have no effect on the concentration of cAMP required for the gene expression to reach its half-saturating concentration. We also showed that the sensitivity of genes depends to a small but significant degree on the differential binding of CRP to the promoters but not RNAP.

### 7. Variation in phenomenological constants across various functional groups

CRP is well known for its role in regulating genes involved in the uptake and utilisation of multiple carbon sources. Apart from this, the cAMP-CRP complex has also been shown to regulate genes involved in nitrogen metabolism, TCA cycle, osmoregulation and antibiotic resistance(34, 41). We binned the genes positively regulated by cAMP into broad functional categories(69, 70). Of these, we focussed on 6 broad categories, namely, Catabolism, Anabolism, Glycolysis, MAF, Respiration, Transport and Stress Response(S7A-B). cAMP regulated genes were enriched for carbon catabolism, transport and respiration (OR > 1, *p-value* < 0.05)(71–73). To see if genes belonging to different functional categories behave differently, we compared the *k, n* and *Emax* of genes across these functional categories (Fig7). We observed no difference in *k* or *n* of genes belonging to any category. However, genes involved in respiration showed significantly higher *Emax* compared to carbon catabolism genes (Wilcoxon rank sum test, corrected *p-value* < 0.01).

**Figure7.**
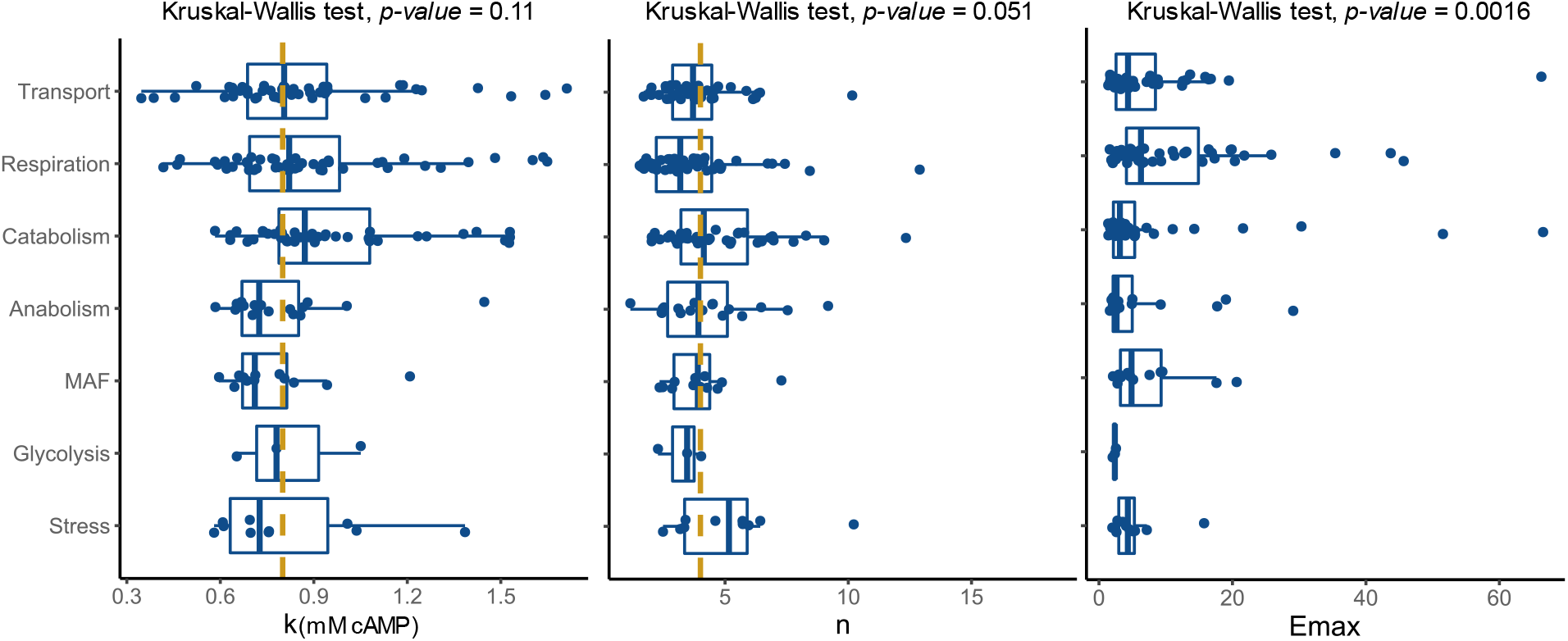
Distribution of k, n and Emax for genes involved in various metabolic pathways. There is no significant differences in *k* and *n* across all metabolic pathways. *Emax* of genes involved in respiration are significantly higher than those involved in catabolism and anabolism isher’s test *adjusted p-value < 0.01*).

Catabolism included genes involved in both uptake and utilisation of various carbon compounds and accounted for 35% of the differentially expressed genes. Most genes in this category were involved in the uptake and breakdown of carbohydrates (67), glycerol (10) and amino acids (13). *E.coli* is able to catabolise a wide range of carbohydrates like simple and complex sugars, organic acids and polyols. In line with this, we observed an upregulation of genes like *fruBKA, malE, malK, lamB, galP, manXYZ, gatYZABCD, mtlAD, uxaB, uxaC, rbsDACBKR, malP* and *malQ. k, n* and *Emax* of carbohydrate genes spanned across the entire biological range of cAMP(S9C-D). Utilisation of a carbohydrate involves both - its uptake via transporters and breakdown by catabolic genes. Many specific and non-specific transporters were activated by cAMP in LB. While the distributions of *k* and *Emax* for catabolic and transport genes do not differ significantly, genes coding for carbohydrate catabolism show significantly higher *n* compared to genes involved in transport (Fig8A-B). This suggests that genes involved in catabolic roles may be under more complex control and behave in a more switch like manner compared to genes coding for transporters. Transporters on the other hand, respond in a graded fashion in response to increasing cAMP dosage.

**Figure8.**
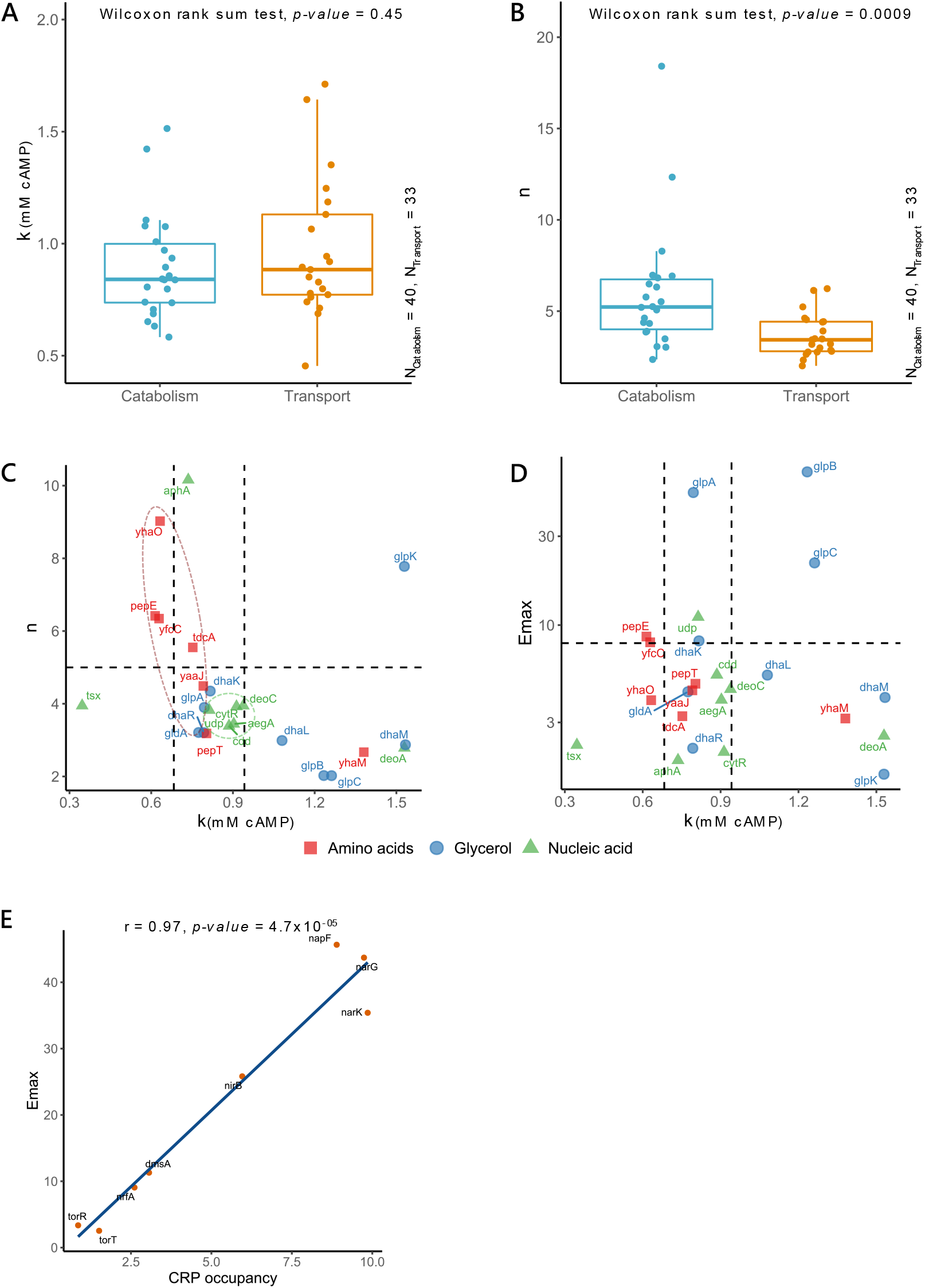
Comparing k, n and Emax across metabolic functional classes. **(A-B)** Distributions of *k* and *n* for carbohydrate catabolism and transport genes. *n* of catabolic genes is higher than those of transport. **(C-D)** Scatterplot showing the relationship between *n* and *k* or *Emax* and *k* for genes involved in catabolism of amino acid, glycerol and nucleic acid. **(E)** Scatterplot showing the correlation between *Emax* and CRP occupancy for anaerobic respiration genes. r and *p-value* represent the Pearson correlation coefficient and the associated p-value for the pair of variables.

*E.coli* can also utilise other molecules as a source of carbon. We observed genes for uptake and utilisation of small peptides and amino acids (*pepE, pepT, sdaC, tdcB* and *tnaA*), nucleic acid derivatives like cytidine and uridine (*cdd, deoA* and *deoC*) and glycerol and phospholipids (*glpABC* and *glpQT*) to express upon addition of cAMP. For this study, we considered values of *n* > 5 to be high, *n* between 3-5 to be moderate and *n* < 3 low. For *k*, we considered values of *k* < 0.68mM cAMP to be low, 0.68-0.94mM to be moderate and *k* > 0.94mM cAMP to be high. We found that most amino acid catabolism genes had low *k* and high *n*, followed by nucleic acid genes which show moderate values for both *k* and *n*, close to the population median. Glycerol and lipid catabolism genes showed moderate to high values of *k* with moderate to low values of *n*(Fig8C-D). This meant that expression of amino acid genes activated at low cAMP concentrations in a switch like manner unlike nucleic acid and glycerol catabolism genes whose response increased in a relatively graded fashion in response to cAMP changes in the cell.

Genes of upper glycolysis (*pfkA, pykA* and *gpmA*) showed an upregulation in response to cAMP. We also found mixed acid fermentation genes (*pflB, adhE, pta, ackA*) to be upregulated in response to cAMP. It is not uncommon for *E.coli* to opt for such overflow metabolism during rapid phases of growth(73, 74). Expression of MAF and glycolytic genes were limited to low and mid ranges of *k* and moderate values of *n*, close to the population median.

We observed an upregulation of respiration genes as well on addition of cAMP. The respiration genes exhibited a significantly higher *Emax* compared to the catabolic genes. Genes involved in respiration spanned the whole range of *k* and *n* and showed very high expression (*Emax*) in response to cAMP. Respiration consisted of genes involved in both aerobic and anaerobic respiration and ETC(S9E-F). Very few aerobic respiration (*cydA, ccmA*) and ETC (*ndh*) genes differentially expressed in LB in response to cAMP. In *E.coli cydA* and *ccmA* encode for cytochromes. These genes responded to low cAMP concentrations in a switch like manner and achieved moderate values of *Emax* ∼5 foldchange. Few anaerobic respiration genes on the other hand exhibited very high *Emax* and spanned across the range of *n* and *k*. These set of genes included *narGHJI, napFDAGHBC* and *dmsBC* genes. We checked the occupancy of CRP at these promoters. We found a strong correlation between *Emax* and CRP occupancy of these genes (Pearson correlation, r = 0.89, *p-value* = 10^−06^, Fig8E). This could explain why genes related to anaerobic respiration express at such high *Emax* despite the cells growing in aerobic conditions.

## Discussion

In this paper we showed the use of phenomenological models like the Hill’s as a tool to gain insights about the cAMP regulatory network. For this study, we generated genome-wide gene expression profiles of *E.coli* as it responds to 10 different concentrations of cAMP. This data revealed the continuous nature of cAMP regulated genes in response to cAMP. Coupled with the use of Hill’s model, we quantify and resolve the 225/305 dose response curves generated in this study. Further, we explored the use of phenomenological constants obtained from the Hills model to understand properties of the cAMP regulatory network.

74% of the cAMP regulated genes followed a sigmoid shaped curve which could successfully be described by the Hill’s model. Our data suggested that promoters and intracellular cAMP concentrations in the cAMP regulatory network may be tuned resulting in optimum gene expression. Most of cAMP regulated genes showed switch-like behaviour with the midpoints of their dynamic range centred around the wild-type concentrations of cAMP (0.72mM cAMP), resulting in a rapid burst of gene expression necessary to reach wild type growth rates in LB as soon as the wild-type concentration is reached. Even after adding concentrations of cAMP much greater (2-4 times) than those found in wild-type cells in LB, the excess cAMP was not able to induce non-LB specific genes. Combinatorial control of environment and transcription factors for gene expression is well known. As cAMP concentrations increase, the cell concurrently unlocks and primes increasing number of metabolic modules. However, environmental signals and metabolic feedbacks tightly control which of these primed modules will actually be expressed in a condition specific manner(21, 17, 19, 75–77, 28).

Expression of cAMP regulated genes differed largely in their *Emax* as compared to *k*. We found that CRP occupancy explained the variation in *Emax* to a small but significant degree, but showed no correlation to *k*. On the other hand, we found gene expression to be independent of the RNAP occupancy at gene promoters. Together these observations imply that – (1) a high affinity CRP promoter need not ensure transcriptional activation at low concentrations of cAMP, instead it is more likely to control the magnitude of a gene’s response to cAMP concentrations and (2) promoter properties other than RNAP and CRP binding alone, play a role in determining the levels of gene expression in response to cAMP. Predictive models for transcriptional regulation as well as some experimental data have shown the contributions of factors like RNAP-CRP synergy, promoter escape rates, gene dosage and presence of secondary co-activators in determining the *Emax*(66–68, 75, 78).

It is well known that feedforward and feedback loops are pervasive in the *E.coli* transcriptional network(49, 61, 79). The presence of high values of *n* for cAMP regulated genes confirmed that. The cAMP regulon showed a lack of genes that respond in a highly switch-like manner(*n*) at high concentrations of cAMP(*k*). This trend remained consistent across both direct and indirect genes(S5D). Based on our null model of a linear regulatory circuit, we expected *n* and *k* of indirect genes to have greater non-linearity and higher variation in their dynamic range. However, observed data showed no differences in the distribution of either *k* or *n* across these set of genes. We suspect that a combination of short length subnetworks, with extensive feedforward and feedback loops feeding into higher nodes could be the reason leading to higher variation in distributions of *k* and *n*. These expectation are based on the fact that a large fraction of cAMP regulated genes are involved in carbon catabolism, which have previously been shown to have short transcriptional circuits with multiple feed-forward loops(80, 79, 81).

In the final section of the paper, we studied how *k, n* and *Emax* are tuned for various metabolic pathways in *E.coli* growing in LB. The most enriched class of genes was carbon catabolism and transport. *k, n* and *Emax* for carbohydrate catabolism and transport genes spanned across the whole measured range. Since genes involved in uptake of different carbohydrates are made up of independent modules, these genes show a broader variation in their *n* and *k*. Contrary to carbohydrates, genes involved in amino acid, nucleic acid and glycerol degradation, which included genes participating in a single pathways, showed more clustered distributions for *n* and *k*(82, 81). However, the number of participating genes were too few to do any statistical analysis on these trends.

We understand that phenomenological models do not have predictive powers and can at best be used as a descriptive tool. Despite the large range of cAMP used in this study, predictions of parameters like *n* and *b0* remained relatively poor owing to lack of resolution at transition points and errors induced by small measures. While it is important to understand the response of genes at a physiological level, this method allowed us to study the response of genes only relative to cAMP and shed very little light on the intricate deeper levels of regulation between individual players in the regulatory circuit. Another caveat is that our study is blind to the composition of ingredients in LB. Thus, for genes that have 2 or more input functions (inducible genes) this approach quantitates only the physiological *k, n* and *Emax*. This makes the values of these phenomenological constants limited to one condition. However, with the advent of cheaper, faster and deeper RNA-seq technologies, such studies can be extended to other carbon sources in a controlled environment.

We find phenomenological models useful to quantitatively describe dose response curves for transcription factors at a global level. This method also circumvents the problems and limitations posed by conventional clustering techniques. Further, we show that the interpretations of the Hill’s parameters can be extended to global regulatory networks to understand its topology. We hope this kind of an approach can provide essential raw material required to ask broader mechanistic questions about transcriptional regulation in bacteria.

## Methods

### 1. Growth conditions and intracellular cAMP measurements

*Escherichia coli* K12 MG1655 was used as the wild-type strain for this study. *ΔcyaA* strain was obtained from Aalap Mogre(37). The *ΔcyaA E.coli* mutant was constructed by deleting the *cyaA* gene from the wild-type strain using methods described in (83). Cells were grown in either Luria-Bertani broth (LB; Hi-Media; catalogue no. M575-500) or M9 minimal media with 0.4% sugars. M9 salts (12.8 gl-1 Na2HPO4.7H2O, 3 gl-1 KH2PO4, 0.5 gl-1 NaCl and 1 gl-1 NH4Cl) were supplemented with 2mM MgSO4, CaCl2, 1mM thiamine, 0.2% Casamino acids and 0.4% (w/v) carbon source (lactose or sorbitol-ribose mixture).

To study the effects of varying cAMP concentrations on growth kinetics and transcriptome of *ΔcyaA E.coli* cells, cAMP sodium salt(Adenosine 3’,5’-cyclic monophosphate sodium salt monohydrate; Sigma-Aldrich; SKU A6885) was added to the growth medium obtaining 10 different final concentrations, ranging from 0mM to 4mM(22). Wild-type *E.coli* cells were used as the control in this experiment. Overnight grown cultures were inoculated at 1:100 dilution in 50ml flasks containing 15ml fresh media with varying amounts of cAMP. Cultures were grown at 37°C and 180 rpm shaking. OD measurements were taken at 600nm every half an hour for the first 4 hours and every 1 hour post that. Maximum growth rate(*µ*_*max*_) of a population was calculated using the growthcurveR package in R(84). The time the bacterial culture took to reach its *µ*_*max*_ was considered to be the “*lag time*”.

Intracellular cAMP concentrations corresponding to the extracellular cAMP was measured using cAMP-ELISA kits provided by Cayman Chemicals. *ΔcyaA* cells grown in different cAMP concentrations, from 0.01mM to 2mM cAMP and cells were harvested when the wild-type population growth reached their *µ*_*max*_. Harvested cells were washed using 1X PBS (8 gl-1 NaCl, 0.2 gl-1 KCl, 1.44 gl-1 Na2HPO4, 0.24 gl-1 KH2PO4 with pH = 7.4) to remove any remaining media and cAMP. Cells were resuspended in 1X PBS and boiled at 95°C for 10 mins. Cell debris was removed by centrifugation at > 10,000rpm(37). The supernatant was analysed using the protocol provided by the cAMP ELISA kits. We found that intracellular cAMP increases linearly with addition of extracellular cAMP(S1G). As a control, intracellular cAMP concentrations of wild-type and *ΔcyaA* mutant were also measured (S1B).

### 2. DNA isolation and sequencing

Genetic backgrounds of wild-type and *ΔcyaA* mutant were confirmed by whole genome sequencing. Overnight grown cultures were used to inoculate 50ml fresh LB media to make a final dilution of 1:100. Cells were grown at 180 rpm at 37°C and harvested at their µmax. Genomic DNA was isolated using GenEluteTM Bacterial Genomic DNA kit (NA2120-1KT, Sigma-Aldrich) using the manufacturer’s protocol. Library preparation was done using Truseq Nano DNA library preparation kits followed by paired end (2×100) sequencing using on Illumina Hiseq 2500 platform. Genome sequences were analysed for SNP and INDELs using the *breseq* software and protocol described by Barrick lab(85). Apart from the expected loss of *cyaA* gene in the *ΔcyaA* mutant, we found a 1063 bp deletion at position 1,977,440 in the *flhC* gene of the *flhDC* operon. This deletion is absent in the wild-type strain. This operon is a known target of the cAMP-CRP signalling system. RNA sequencing results show that the *flhDC* operon and downstream gene *fliA* encoding for σ^F^ fail to respond to high doses of extracellular cAMP resulting in shutting down of flagellar, motility, chemotaxis and biofilm related genes in the *ΔcyaA* mutant(86, 87).

### 3. RNA sequencing and differential expression analysis

*ΔcyaA* cells grown in LB with different cAMP concentrations were harvested when the wild-type population reached *µ*_*max*_. Two replicates for each sample were processed. Total RNA was isolated using TRIzol-chloroform extraction method, followed by DNase treatment. 16S and 23S ribosomal RNA were depleted using Ambion MICROBExpress bacterial mRNA enrichment kits (AM1905) and the RNA quality checked using Bioanalyzer (Agilent) followed by Qubit quantification. Libraries for each sample were prepared using the New England Biolabs (NEB) NextUltra directional RNA library prep kit followed by single end sequencing using Illumina HiSeq 2500 platform.

All annotation and sequence files were obtained from NCBI. *E.coli K*-12 MG1655 (NC_000913.2) was used as the reference genome for RNA sequence analysis. Sequencing reads were aligned and mapped to the reference genome using the Burrows-Wheeler Aligner (BWA) algorithm. SAMtools (v1.2) and BEDtools (v2.25.0) were used to determine read counts per gene. Normalisation and differential gene expression analysis across samples was done using the EdgeR package(3.28.1) in R as described by Chen et al(88).

For any pairwise comparison, differentially expressed genes were defined as the set of genes showing a logFC ±1 and *p-value* < 0.01. Genes differentially expressed between the wild-type strain compared to the *ΔcyaA* mutant were considered to be cAMP responsive. Effect of any cAMP concentration on a particular gene was quantified as the foldchange a gene experienced at that cAMP concentration compared to the 0mM cAMP *ΔcyaA* state. Since log scale is non-linear and will affect the shape and magnitude of individual trends, all log foldchange values produced by EdgeR were converted to foldchange before further analysis.

We find that genes coding for flagella, chemotaxis and biofilm formation, which are under the control of the *flhDC* and σ^F^ do not respond to even high concentrations of extracellular cAMP. For this study we have removed them from the set of cAMP responsive genes(S2A).

### 4. Chromatin Immunoprecipitation (ChIP-Seq) for CRP

ChIP method was adapted from a previous study with few changes(59). Cells were grown aerobically at 37°C to the early exponential phase (∼O.D. 0.2). Formaldehyde was added at a final concentration of 1% and incubated for 20 minutes at 37°C. Glycine was added to quench the cross-linking at a final concentration of 0.5M and cells were incubated for 5 minutes. Cells were then harvested and washed three times with cold 1X TBS buffer and resuspended in 1 ml of lysis buffer (10 mM Tris (pH 8.0), 20% sucrose, 50mM NaCl, 10mM EDTA, 20 mg/ml lysozyme and 0.1 mg/ml RNase A) and incubated at 37° C for 30 minutes. After the incubation, 3ml of Immunoprecipitation buffer (IP buffer) [50 mM HEPES–KOH (pH 7.5), 150mM NaCl, 1mM EDTA, 1% Triton X-100, 0.1% sodium deoxycholate, 0.1% sodium dodecyl sulphate (SDS) and PMSF (final concentration 1 mM)] was added. Cells were then sheared in the Bioruptor (Diagenode) with 33 cycles (25 seconds on/24 seconds off). Cellular debris was removed by centrifugation at 4°C and the supernatant was split into aliquots for ChIP and input samples. Each aliquot was incubated with 1X TBS pre-washed 20 μl of protein A/G ultra-link resin beads on a rotary shaker for 45 minutes at room temperature. Samples were then centrifuged and the supernatant was incubated with flag mouse monoclonal antibody (Sigma-Aldrich) (IP), and no antibody (Mock IP) for 60 minutes at room temperature. Meanwhile, 40 μl A/G ultra-link resin beads (ThermoFisher Scientific) per sample were blocked in 1 mg/ml BSA (Bovine Serum Albumin). After one hour of incubation, 40 μl blocked A/G ultra-link resin beads were added to all samples and incubated for 90 minutes at room temperature. Beads were then collected after centrifugation in SpinX-Costar tubes. After collection, beads were then washed successively in IP buffer, twice with High Salt IP buffer (IP buffer + 500 mM NaCl), once with wash buffer [10 mM Tris (pH 8.0), 250mM LiCl, 1mM EDTA, 0.5% Nonidet P-40 and 0.5% sodium deoxycholate] and once with TE [Tris-EDTA (pH 7.5)]. All samples were washed rotating the tubes in rotary shaker for 3 minutes, and centrifugation at 3500 rpm for 2 minutes. Immunoprecipitated complexes were eluted in 100 μl elution buffer [10 mM Tris (pH 7.5), 10mM EDTA and 1% SDS at 65° C for 20 minutes. After elution, the sample along with the input was reverse cross-linked in 0.5X elution buffer + 0.8 mg/ml Pronase (Sigma-Aldrich) at 42° C for 2 hours followed by 65° C for 6 hours. DNA was then purified using Minelute PCR purification kit (Qiagen). After ChIP experiment, 5 ng of DNA from antibody treated and input was taken to prepare library using Illumina TruSeq DNA preparation kit. All the steps were performed using manufacturer’s instructions and proceeded with paired end sequencing. The paired end reads after adaptor trimming were aligned to the E. coli reference genome (NC_000913.2) using BWA. Reads per base was calculated using SAMtools(89).

### 5. Estimating CRP and RNAP occupancy at gene promoter

In-vivo CRP and RNA polymerase occupancy were calculated from CRP and RNAP ChIP-seq experiments performed in *E.coli* cells at an exponential phase in LB. The occupancy at any genomic region was considered to be proportional to the intensity of the ChIP signal at that base pair. Occupancy, in turn, is used as a proxy for the affinity of CRP and RNAP for that region.

To calculate occupancy at specific genomic regions from ChIP-seq data, reads per base pair were obtained. It was ensured that the reference genome used for alignment in ChIP studies corresponded to the one used for RNA sequencing experiments (NC_000913.2). For both data sets, reads per base were internally normalised by dividing the frequency at each base by the mode of the distribution and the signal at each base was calculated as the ratio of the test signal to that of the internal control. Spurious peaks were removed by smoothing the data using local regression method. The maximum frequency recorded within a given region was considered as the occupancy for that region. A region of -200 to +50 base pairs flanking the start site of the gene was considered the CRP or RNAP binding region. For genes which are part of operons, the occupancy score for only the first gene was considered for all analyses.

CRP affects gene expression by recruitment of RNAP at the promoter. Despite being from two different studies, a significant correlation between CRP and RNAP occupancy (Pearson correlation coefficient, r = 0.36; p-value = 7.7 × 10-06; S6E) was observed, showing that the data sets are internally consistent.

### 6. Model fitting and estimation of parameters

All dose-response curves were calculated as the foldchange of a gene at a given cAMP concentration compared to the *ΔcyaA* mutant. All analysis was done using custom scripts in R. All curves were fit to different models using the *nls* function from R stats package.

#### a) Model definition

A gene is expressed when the cAMP-CRP active complex binds to the promoter of a gene. Based on the underlying regulatory mechanisms, gene expression versus cAMP concentration response curves may follow one of the trends – no response, linear/non-saturating or sigmoid. We checked the fit of each gene to these 3 models.

### Hill’s Model (HM)

Non-linear relationships, especially sigmoidal ones are common in transcriptional control of gene expression by activators and inhibitors(60). Gene expression following a sigmoidal response curve can be captured well using phenomenological models like the 4 parameters Hill’s model, defined by *b0, n, k* and *Emax*. Hill’s model can be derived by considering the equilibrium binding of a transcription factor to its promoter(90, 91, 51, 53, 57). We held the following assumptions while applying the Hill’s model to our data - (1) the extracellular cAMP immediately equilibrates with the intracellular cAMP; (2) the active cAMP-CRP complex is proportional to extracellular cAMP and (3)RNAP is a not a limiting factor for transcription(92). For an activator, like the cAMP-CRP complex, the Hill’s curve can be written as:

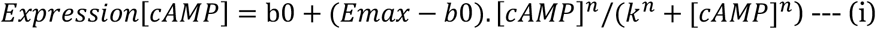

The biological relevance of the phenomenological constants is well hashed out in the field(50, 56). We extend these definitions to a network:

1. *bo* is the baseline expression of a gene in the cell when cAMP is absent in the system. Biologically, this can be interpreted as the leaky expression of the system and depends on how tightly repressed a gene is. In our study, *b0* is ∼1 foldchange.
2. *Emax* represents the expression level when the concentration of the cAMP-CRP complex has far exceeded its binding sites and transcription is no longer limited by the concentration of intracellular cAMP. Physically, it is affected by intrinsic promoter properties like RNA polymerase (RNAP) binding strength to the promoter, the interaction of cAMP-CRP or other NAPs binding at the promoter, interaction between the cAMP-CRP complex and RNAP, effects of gene dosage and promoter escape rates on levels of transcription(67, 68, 78, 93). Biologically, *Emax* tells us the maximum sensitivity of a gene in response to cAMP. *Emax* is measured in terms of foldchange compared to the *ΔcyaA* mutant and ranges from 1 to infinity.
3. *k* represents the cAMP concentration at which half the saturating expression has been achieved. Biologically, it is the midpoint of the dynamic range of the gene. For switch-like genes, it reflects the cAMP concentration at which the gene starts expressing. Physically, for genes which are regulated by the direct binding of the cAMP-CRP complex to their promoters, *k* reflects the affinity of the complex to the promoter. For genes that are under indirect regulation of cAMP, *k* reflects the composite effects of all *k*s in the network(57).
4. *n* determines the extent of graded versus switch type response the gene has upon activation. For a single promoter-cAMP-CRP complex pair, this reflects the cooperative behaviour of the transcription factor. It implies, either enhanced binding or decreased unbinding of the transcription factor, as a function of cAMP concentration. For gene regulatory networks, *n* indicates the presence of positive feedback and multistep feedforward loops(60, 62, 91). Steeper the response curve, higher is the *n*.

### Linear/Non-saturating Model (LM)

Genes whose saturating concentrations are well beyond the cAMP concentrations administered in our study will fail to show saturation under our cAMP regime. The Hill’s equation from the previous section can be modified to a non-saturating model as follows:

For *k* >> [cAMP];

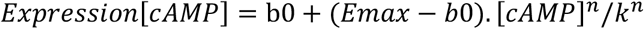

Since, (*Emax-bo)/k*^*n*^ will be a constant for each given curve, the above equation can be reduced to:

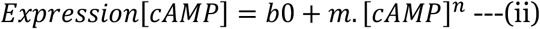

A linear relationship between gene expression and cAMP concentration is a special case of (ii) where n = 1. This can mathematically be represented by:

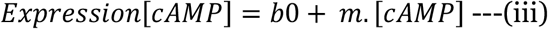

where *m* is the slope of the line and *bo* the basal expression of the gene.

### Non-responsive Model (NR)

The null model for gene expression response to cAMP is given by a no expression model:

For [cAMP] = 0

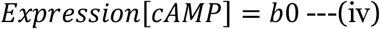

#### b) Goodness of fit measurements

The best fit for each model was estimated using methods that minimise the sum of squared estimate of errors (SSE), where the SSE is defined by:

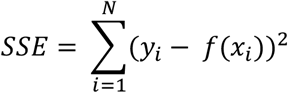

where, for *N* number of observations, *y*_*i*_ is the i^th^ value of the variable to be predicted, *x*_*i*_ is the i^th^ value of the explanatory variable and *f(x*_*i*_) is the predicted value of *y*_*i*_.

A model with lower estimates of SSE was considered to be the better fit. SSE is a meaningful measure when comparing competing models. However, it does not reflect on how well the fitted model explains the observed data. For a linear regression, the R^2^ coefficient of determination gives a statistical measure of how well the regression predictions explain the observed data. R^2^ is measured as:

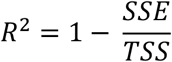

where TSS is given by:

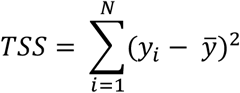

where, *y*_*i*_ is the i^th^ value in the sample and 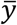 is the mean of the sample.

However, R^2^ is not calculated for non-linear curve fits and has been shown to give inconsistent results. Here, we try to define a similar metric for the non-linear regression model. To do so, a R^2^ value was computed between the observed values and the values predicted using the fitted function for each cAMP concentration. This R^2^ value obtained quantified how well the variation in the observed value is explained by the model fitted predicted data. Along with R^2^ values, the slope between the observed data and predicted data was also calculated.

Finally, to determine the accuracy of each predicted parameter – *bo, n, k, Emax* or *m*, a relative standard error (RSE) was calculated by:

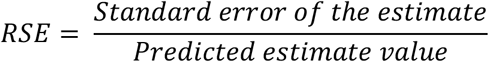

#### c) Model fitting and parameter estimation

The *nls* function from the R Stats package was used to fit the three models-no response, non-saturating/linear and Hill, to each gene curve. The *nls* function returns the residual standard error (*σ*^*2*^) value for the best possible fit given each model.

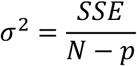

where *N* is the number of observations and *p* is the number of parameters in the model.

The estimated parameter values and their standard errors extracted from the fit were used to calculate the corresponding RSE for the predicted parameters. The *predict.nls* function was used to predict expression levels from the fitted function at each cAMP concentration. R^2^ values were calculated using custom codes as defined in the previous section.

A gene was considered to behave sigmoidal only if it’s *σ*^*2*^ for the Hill’s model (equation i) fit was less than that of the other models, R^2^ > 0.80 and RSE for *Emax* < 20% and *k* < 20%. Since no genes are differentially expressed between 0.01mM cAMP and *ΔcyaA*, all genes have a *bo* in the range 0.8-1.2 foldchange. Since the number of data points around the transition states is few, estimations of *n* showed higher standard errors compared to *k* and *Emax*. Hence, for this study RSE of the estimated *bo* and *n* are not used as filters to determine sigmoid genes. 225/305 genes satisfied these criteria.

A gene was considered non-saturating, if the *σ*^*2*^ for the fit for Non-saturating model (equation ii) was lower than any of the other models and its R^2^ > 0.7 with a *p-value* < 0.05. Similarly for the Linear model (equation iii). 28/305 genes satisfied both the models. Very small difference in the RSE as well as R^2^ values for genes that fit both Non-saturating and Linear models were found. Thus, for the rest of the paper, this group is referred to as Linear Model.

Finally, genes were considered to be non-responsive if the *σ*^*2*^ for the fit to Non-responsive model (equation iv) was less than that of any other models and R^2^ > 0.7 and *p-value* < 0.05. Genes that did not fit any of the categories were analysed manually. Out of the 305 DE genes, 225 genes were found to follow Hill’s model (HM), 28 genes were best explained by a linear model (LM) and 18 genes failed to respond to extracellular cAMP (NR). We found 34 genes to follow a non-monotonic inverted U trend in response to increasing cAMP concentrations (NM).

### 7. Cluster analysis

In order to cluster the genes based on their trends, dose-response curves of each cAMP regulated genes were Z-score normalised and then partitioned using conventional clustering methods like *k-means* and *Hierarchical* clustering. Hierarchical clustering was performed using the *hclust* function from the R Stats package using Pearson distance as the dissimilarity measure and average linkage as the clustering method. The pheatmap package from R was used for the visualisation of clustered heatmaps. We wanted to validate the use of Pearson distance as a useful metric for clustering gene trends using hierarchical clustering. Genes belonging to the same operon (Intra-operonic genes) should show high correlation values between the trends they follow compared to non-operonic genes (Inter-operonic). We used this comparison as an internal control. Intra-operonic genes showed a higher and tighter spread of Pearson correlation values compared to inter-operonic genes(S3D), validating that Pearson distance can be used as a metric to cluster our data set.

For *k*-means clustering, the optimum number of clusters was determined using WSS, Gap-stat and Silhouette methods. The *kmeans* function from the base R Stats package was used to divide the genes into the desired number of clusters. A PCA plot was used to visualise the clusters formed. *prcomp* from the R Stats package was used to calculate the principal components and relative position of each gene.

### 8. Data Availability

RNA-seq data is available at GEO with the accession GSE202549. ChIP-seq data is available at GEO with accession number GSE104505.

All codes are available at https://github.com/shwetac09/cAMP_project_codes_2022.git

