## Supplementary figures for "Understanding the genome-wide transcription response to varying cAMP levels using phenomenological models in bacteria"

### Supplementary 1

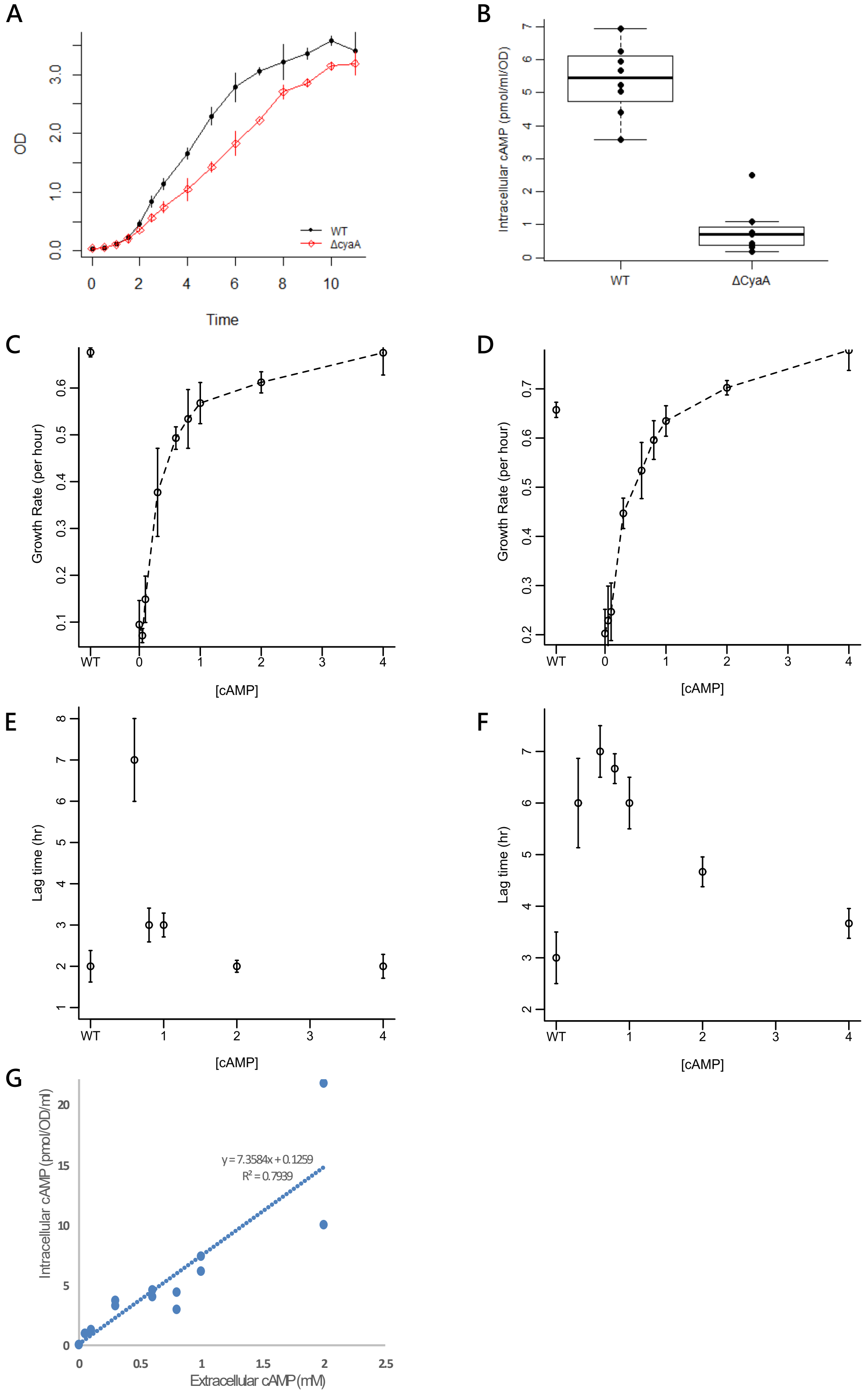


**S1.(A)** Growth curve of wild-type and *ΔcyaA E.coli* cells in LB. Growth rates for these strains were calculated using these growth curves.**(B)** Intracellular cAMP concentration in wild-type and *ΔcyaA* strain in LB. **(C-D)** Response of growth rates to increasing doses of cAMP concentration for *ΔcyaA* cells when grown in M9 sorbitol-ribose and M9 lactose media respectively. **(E-F)** Response of lag times to increasing doses of cAMP for *ΔcyaA* mutants growing in M9 lactose media and M9 sorbitol+ribose respectively.**(G)** Relationship between intracellular and extracellular cAMP concentrations.

### Supplementary 2

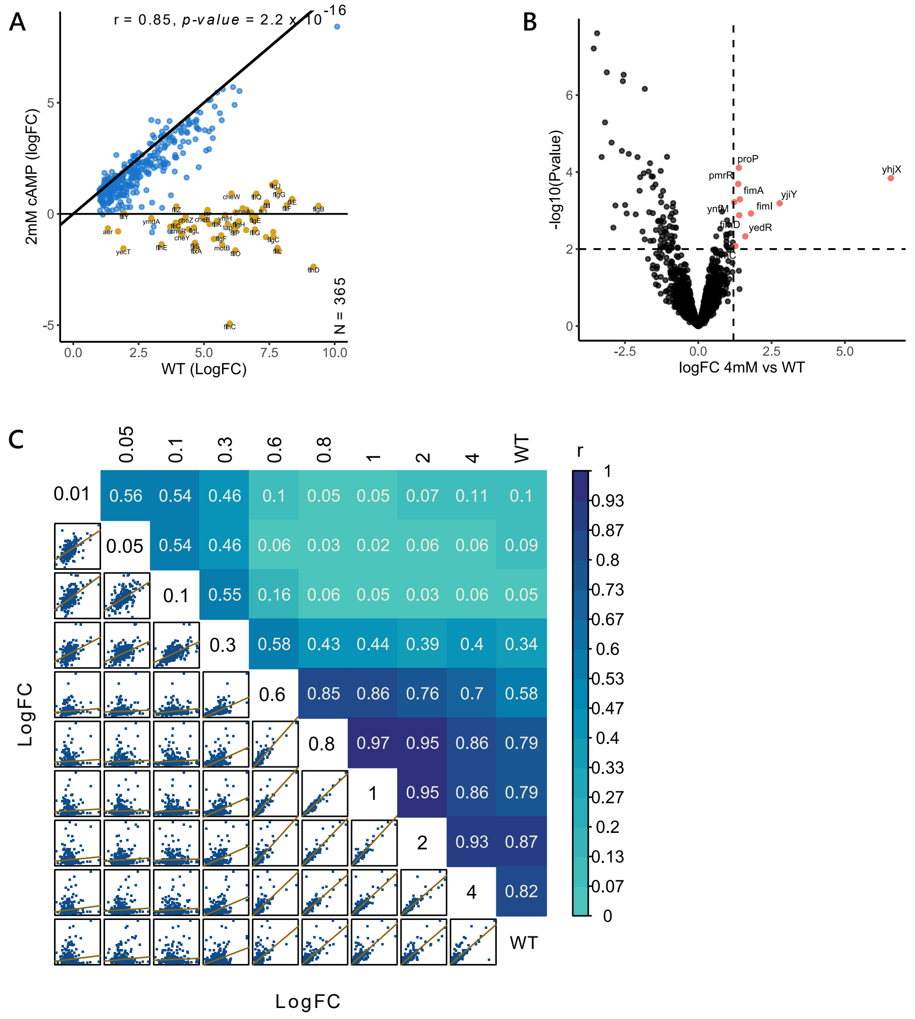


**S2. (A)** Scatterplot showing the correlation between the gene expression at 2mM cAMP and wild-type (WT). Despite addition of excess extracellular cAMP, few genes failed to respond to cAMP (yellow). **(B)** Correlation plot showing pairwise comparison of transcriptome of at each cAMP concentration. Shown are the scatterplots with trendline for all pairs of comparison (lower half) and the corresponding Pearson correlation coefficient (upper half).

### Supplementary 3

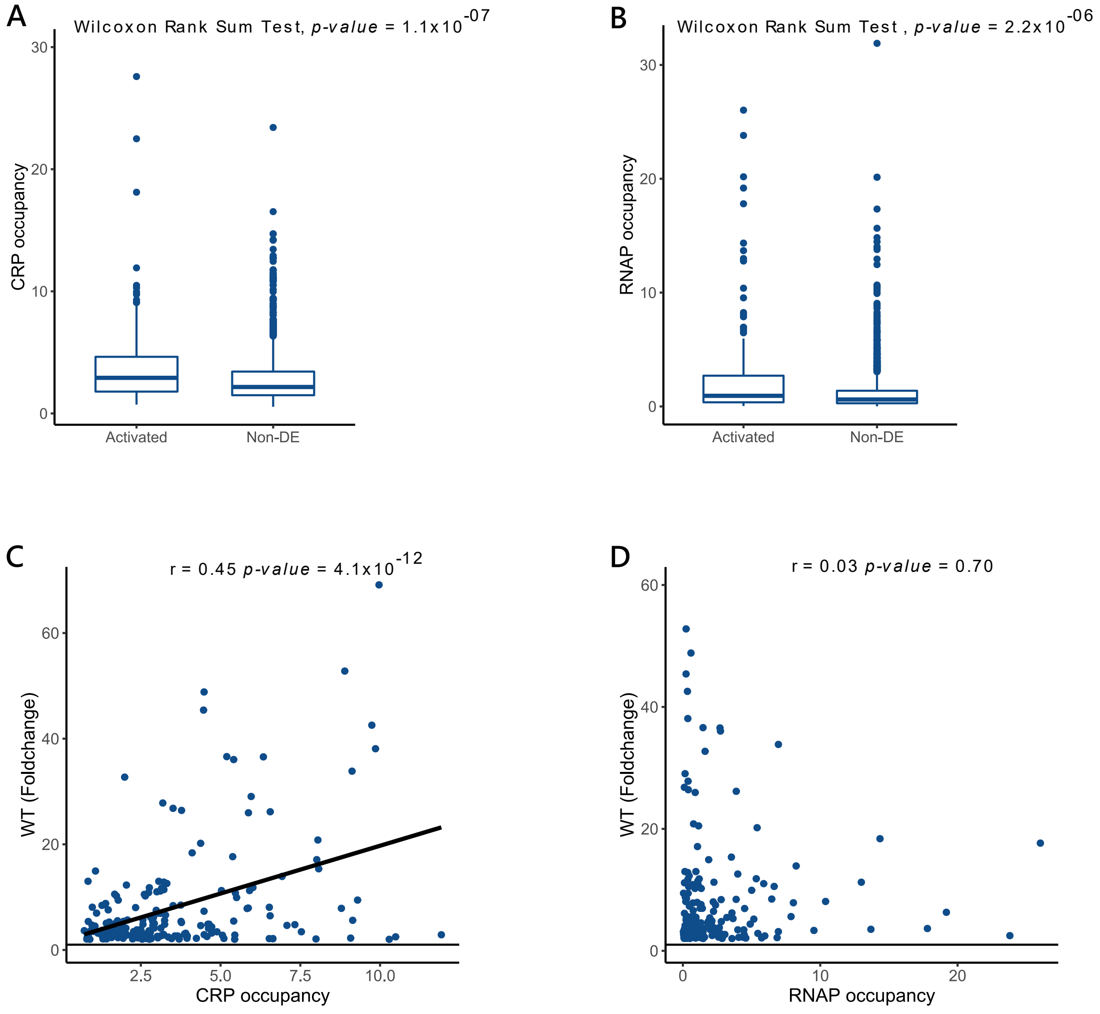


**S3. (A-B)** CRP occupancy and RNA polymerase occupancy scores at promoter for genes which are positively regulated by cAMP and genes which are not differentially expressed. As expected, CRP occupancy and RNA polymerase occupancy at transcribing genes is higher than non-transcribing genes. **(C-D)** Correlation between wild-type levels of gene expression compared to delta-*cyaA* (in foldchange) and CRP occupancy(C) and RNAP occupancy(D) at the promoter of cAMP regulated genes.
r and *p-value* indicate the Pearson correlation coefficient and associated *p-value* for the give pair of variables.

**Supplementary 4**

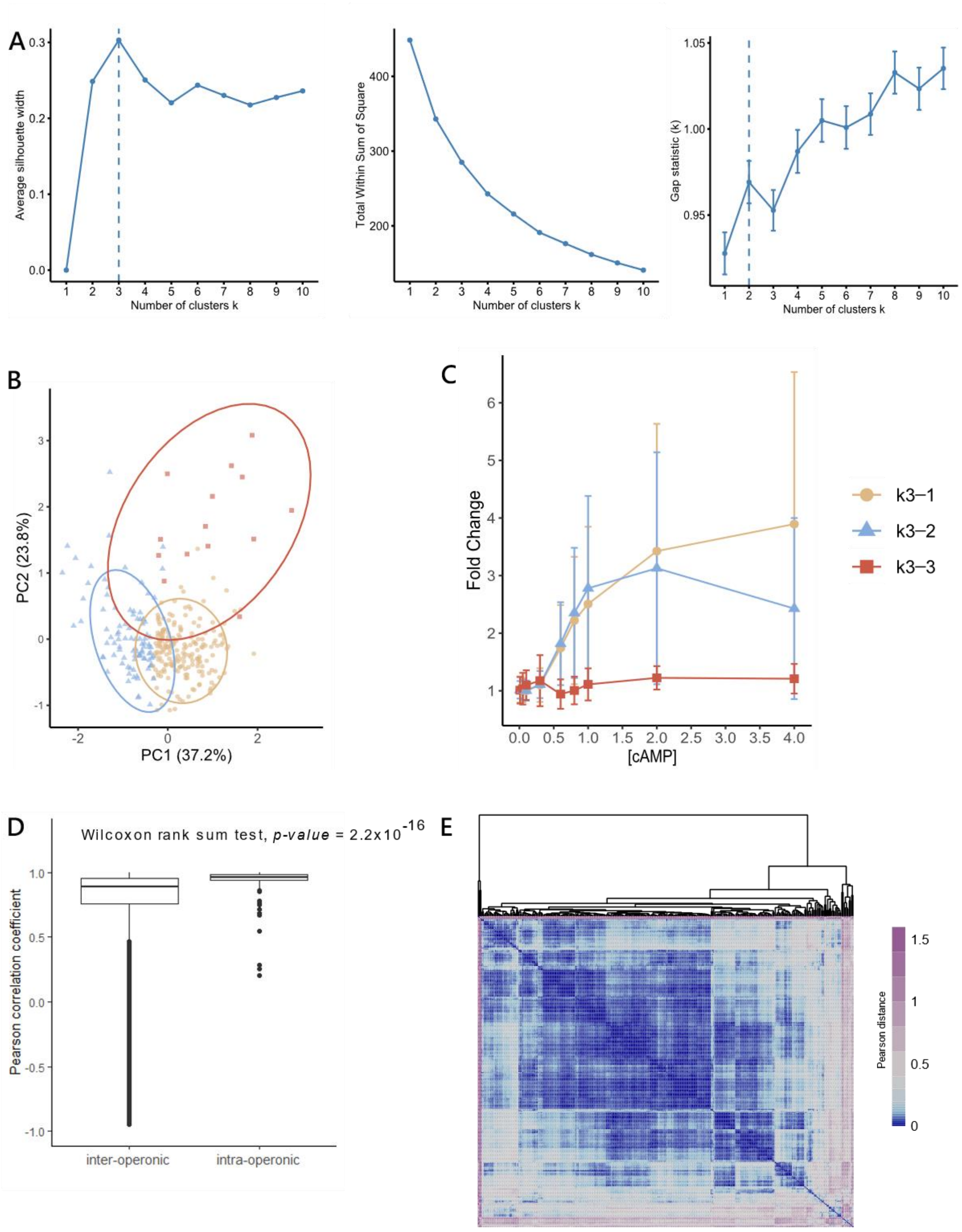


**S4. (A)** Optimum clusters for k-means using 3 different method – Silhouette, WSS and Gap-statistic. There is no consensus in the number of predicted clusters. We divide the data into 3 clusters based on the Silhouette method.**(B)** PCA plot showing the 3 clusters formed by k-means algorithm. It shows the continuum in gene expression trends varying smoothly along PC1. **(C)** Median trends for the 3 clusters obtained by the k-means clustering algorithm. Error bars represent the median absolute deviation. **(D)** Boxplot showing the distributions of Pearson correlation coefficients between inter and intra-operonic genes. This plot shows that operonic genes have trends which are more tightly correlated than genes belonging to different transcription units. Thus, Pearson correlation coefficient can be considered a suitable distance metric for clustering genes. **(E)** Dendrogram generated by hierarchical clustering of expression trends of cAMP regulated genes using Pearson correlation coefficient as the distance metric and average as the linkage method. Heatmap shows the Pearson distance between genes, with blue indicating higher correlations and violet indicating low correlation values.

### Supplementary 5

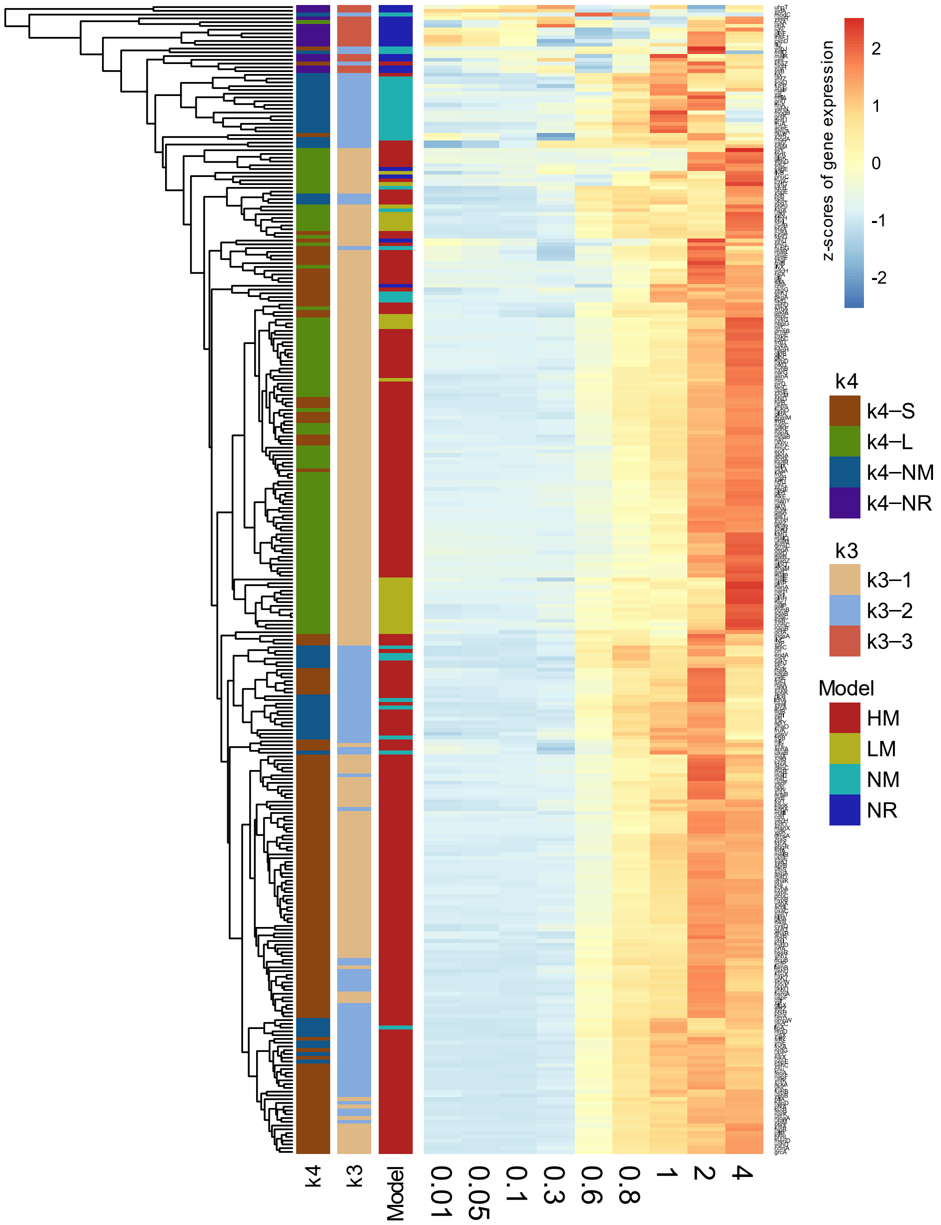


**S5**.Comparison of all 3 clustering methods – hierarchical, k-means and model-fit. Each row represents a gene. The heatmap shows normalised gene expression in response to the indicated cAMP concentration. Broadly gene cluster similarly across all the methods. Clusters from hierarchical and k-means clustering algorithms are in agreement with each other to a greater degree compared to the model fit method.

### Supplementary 6

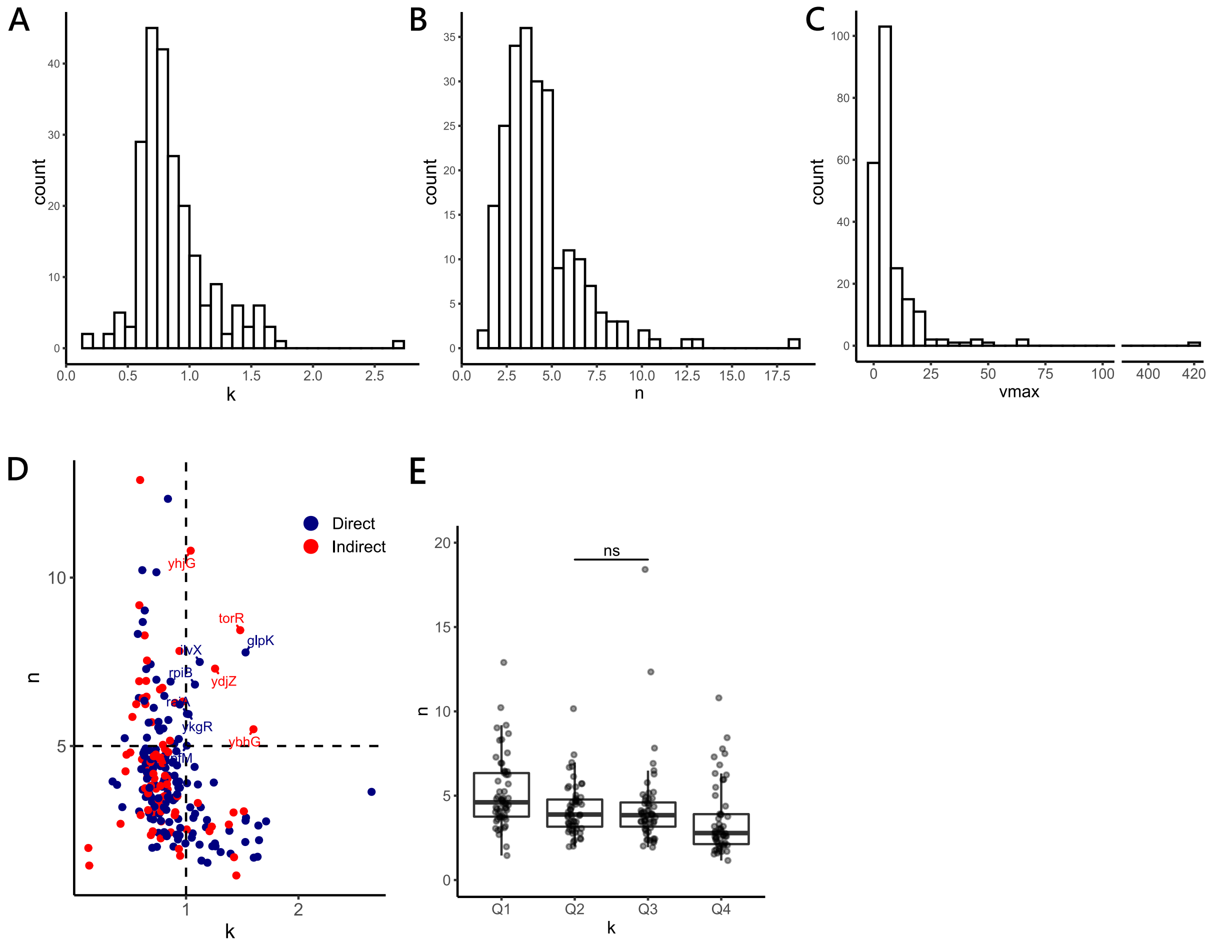


**S6. (A-C)** Histograms for *k*, *n* and *Emax* values for cAMP regulated genes. **(D)** Distribution of *n* versus *k*, with coloured dots representing genes under $Direct_{ChIP}$ (blue) and $Indirect_{ChIP}$ (red) control of cAMP. **(E)** Boxplot showing that genes with lower *k* have significantly higher *n* compared to genes with higher *k*. Q1 to Q4 represent the genes belonging to 4 quartiles of *k*. Except for Q1 and Q2, all other combinations have significantly different distributions of *n* (Wilcoxon rank sum test, p-value < 0.01).

### Supplementary 7

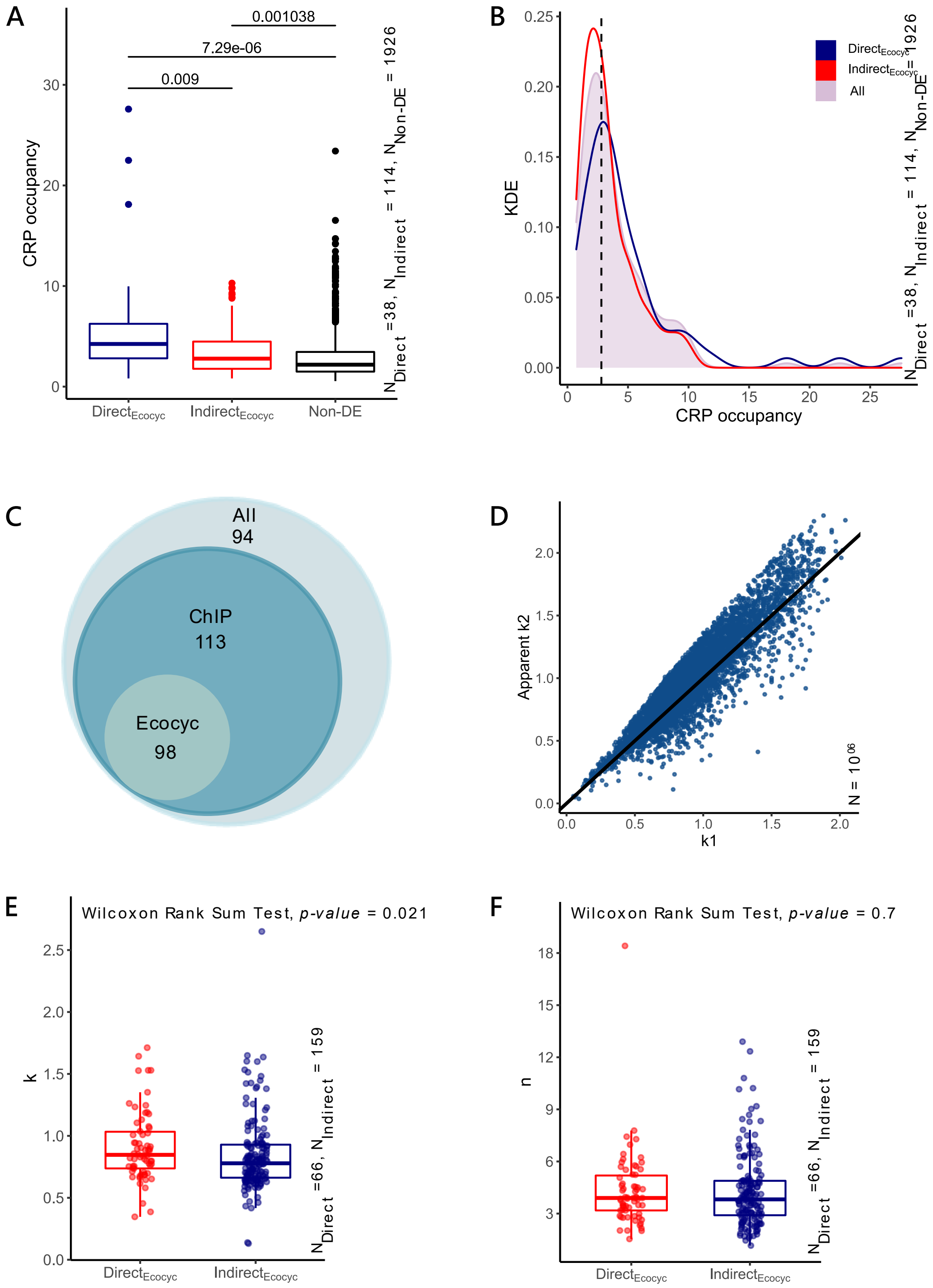


**S7. (A)** cAMP regulated genes are binned as direct or indirect targets of cAMP based on data from the Ecocyc database. Comparison of CRP occupancy at promoters of $Direct_{Ecocyc}$ and $Indirect_{Ecocyc}$ genes with genes not differentially expressed(Non-DE) under the cAMP regime. Unexpectedly, $Indirect_{Ecocyc}$ genes show greater CRP occupancy compared to non-DE genes, suggesting presence of unidentified directly regulated genes in the $Indirect_{Ecocyc}$ set. **(B)** Kernel density distribution of CRP occupancy scores for set of $Direct_{Ecocyc}$ and $Indirect_{Ecocyc}$ genes, with the dashed line showing the Q1 (bottom first quartile) of the CRP occupancy score of the $Direct_{Ecocyc}$ genes. Q1 = 2.8. This is used as the cut-off for determining a broader set of direct genes - $Direct_{ChIP}$ gene set. **(C)** Venn diagram showing direct genes as defined by Ecocyc and ChIP cut-off as a subset of all cAMP regulated genes. **(D)** For a simple linear regulatory network toy model, scatter plot showing the spread of apparent *k2* relative to *k1*. Apparent *k2* can be less than *k1*. **(E-F)** Distributions of *k* and *n* across $Direct_{Ecocyc}$ and $Indirect_{Ecocyc}$ genes.

### Supplementary 8

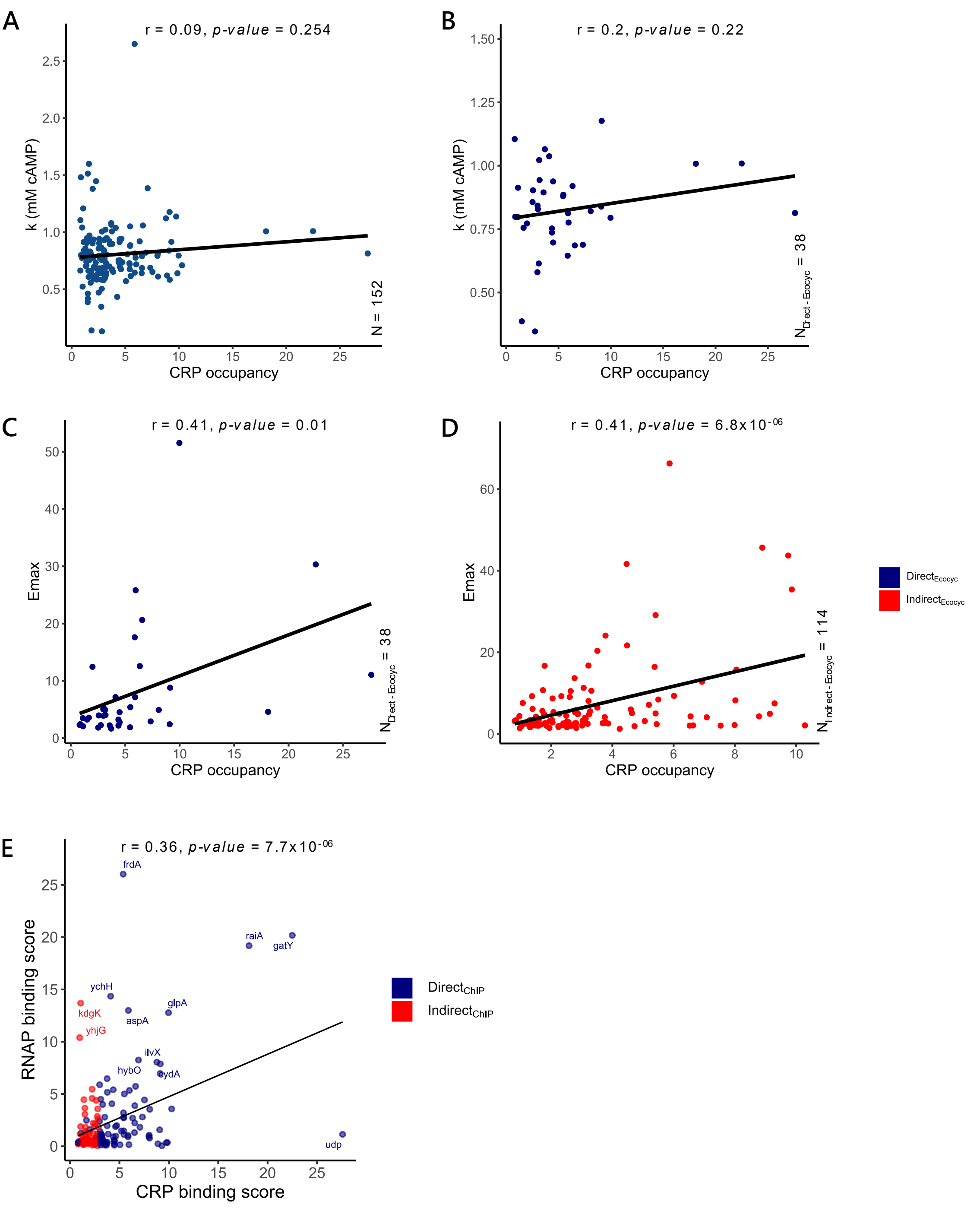


**S8. (A-B)** Scatter plot showing variation of k with CRP occupancy across all cAMP regulated genes (A) and for the $\text{Direct}_{\text{Ecocyc}}$ set of genes(B). Throughout the figure, blue colour represent genes under direct regulation of cAMP and red represent indirect genes. Subscripts show which method of binning was used to determine the direct and indirect set of genes.**(C-D)** Variation of Emax with CRP occupancy of $\text{Direct}_{\text{Ecocyc}}$ and $\text{Indirect}_{\text{Ecocyc}}$. Contrary to expectations, we observed a moderate correlation between Emax and CRP occupancy of $\text{Indirect}_{\text{Ecocy}}$ genes. However, $\text{Direct}_{\text{Ecocyc}}$ genes show a positive correlation between Emax and CRP occupancy. **(E)** Scatterplot showing the positive correlation between RNAP occupancy scores and CRP occupancy scores of across all cAMP regulated genes..
r and p-value indicate the Pearson correlation coefficient and associated p-value for the give pair of variables.

### Supplementary 9

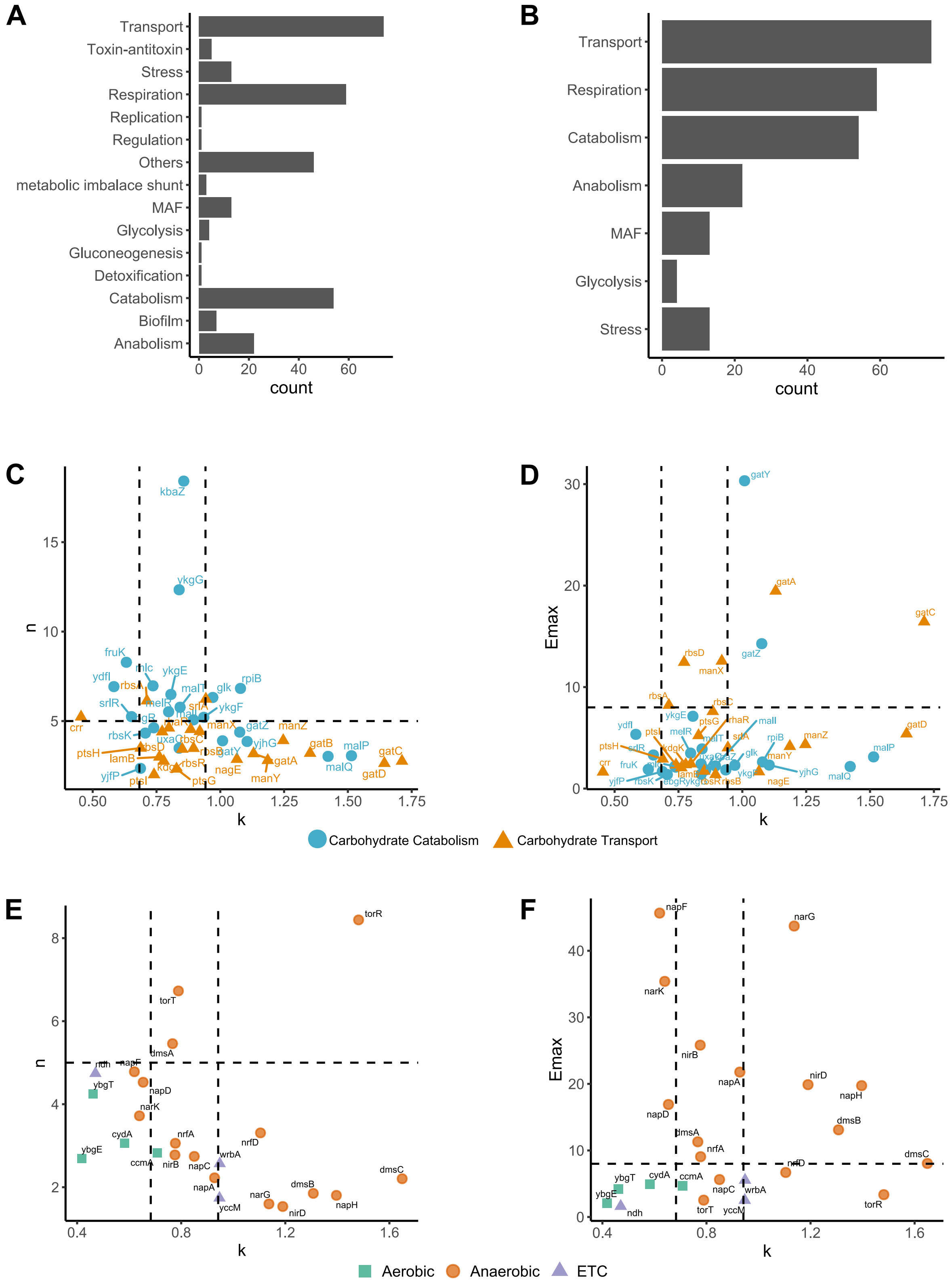


**S9.(A-B)** Counts of genes belonging to different functional categories – (A) all categories that Multifun/Ecocyc binned cAMP regulated genes into. (B) Categories of interest in this study.**(C-D)** Scatterplot for *n* versus *k* and *Emax* versus *k* for carbohydrate genes. Different colours represent genes associated with catabolism (blue) or transport (yellow). **(E-F)** Scatterplot for *n* versus *k* and *Emax* and *k* for genes involved in respiration. Different colours/shapes represent the type of respiration (Aerobic/Anaerobic) and genes involved in forming the ETC complex in *E.coli*.
